# Widespread inhibition, antagonism, and synergy in mouse olfactory sensory neurons *in vivo*

**DOI:** 10.1101/803908

**Authors:** Shigenori Inagaki, Ryo Iwata, Masakazu Iwamoto, Takeshi Imai

## Abstract

Sensory information is selectively or non-selectively inhibited and enhanced in the brain, but it remains unclear whether this occurs commonly at the peripheral stage. Here, we performed two-photon calcium imaging of mouse olfactory sensory neurons (OSNs) *in vivo* and found that odors produce not only excitatory but also inhibitory responses at their axon terminals. The inhibitory responses remained in mutant mice, in which all possible sources of presynaptic lateral inhibition were eliminated. Direct imaging of the olfactory epithelium revealed widespread inhibitory responses at OSN somata. The inhibition was in part due to inverse agonism toward the odorant receptor. We also found that responses to odor mixtures are often suppressed or enhanced in OSNs: Antagonism was dominant at higher odor concentrations, whereas synergy was more prominent at lower odor concentrations. Thus, odor responses are extensively tuned by inhibition, antagonism, and synergy, at the early peripheral stage, contributing to robust odor representations.

## INTRODUCTION

Sensory systems have to detect biologically meaningful sensory stimuli from background signals. Various types of inhibition play important roles in this purpose. For example, the visual system has to detect a visual cue under ambient light. In order to detect specific visual features, both decreases and increases in neurotransmission from photoreceptor cells are conveyed to the ON and OFF pathways, respectively. In addition, lateral inhibition within the retinal circuits enables the spatial contrast enhancement, thus aiding the detection of a visual object(Demb and Singer, 2015).

Similarly, the olfactory system has to reliably detect odorants under various background signals. Natural environments often contain a lot of ambient odorants. Also, accumulating evidence shows that many OSNs robustly respond to mechanical stimuli produced by the nasal airflow (Chen et al., 2012; Connelly et al., 2015; Grosmaitre et al., 2007; Iwata et al., 2017). As a result, sniffing alone produces responses in many OSNs without odors. Nevertheless, animals can often identify a component from a mixture of odorants (Rokni et al., 2014). How specific odor information is reliably extracted from the background signals is a central issue in the field. The olfactory bulb (OB) and cortical circuits certainly play important roles in this task (Imai, 2014; Wilson and Mainen, 2006). In particular, the roles of inhibitory circuits have been well recognized, in terms of gain control, lateral inhibition, and temporal patterning (Banerjee et al., 2015; Economo et al., 2016; Fukunaga et al., 2014; Kato et al., 2013; McGann et al., 2005; Yokoi et al., 1995). However, we know little about odor information processing at the entry point of the olfactory system, the OSNs, particularly in the physiological conditions *in vivo*.

In the mammalian olfactory system, each OSN expresses just one type of odorant receptor (OR) out of a large repertoire (~1,000 in mice), and OSNs expressing the same type of OR converge their axons to a set of common glomeruli in the OB. It is also known that all types of ORs in the main olfactory system couple to G_olf_, and cAMP signals induce depolarization of OSNs via CNG channels (Firestein, 2001). So far, most electrophysiological or calcium imaging studies of OSN somata have been limited to isolated OSNs or olfactory epithelium (OE). Therefore, it has been difficult to obtain comprehensive odor response profiles at the level of OSNs. Based on our limited knowledge, it has been generally believed that odorants “activate” a specific set of ORs (known as the “combinatorial receptor code”) (Malnic et al., 1999; Saito et al., 2009), and odor information is represented by the map of “activated” glomeruli in the OB (Mori et al., 2006).

Here we performed GCaMP calcium imaging of OSNs *in vivo* and found widespread inhibitory responses at OSN axon terminals in the OB. We also found that inhibitory responses are already evident at the OSN somata. Moreover, a response to an odor mixture demonstrated extensive suppression (antagonism) as well as a supra-linear enhancement (synergy), contrary to the linear summation model. Thus, inhibition, antagonism, and synergy at the receptor level tunes odor responses in the mammalian olfactory sensory neurons.

## RESULTS

### Inhibitory responses at OSN axon terminals *in vivo*

Characterizing odor responses at the most peripheral level is fundamental to our understanding of odor information processing in the mammalian olfactory system. In this study, we used an OSN-specific GCaMP transgenic mouse line, OSN-GCaMP3 (*OMP-tTA; TRE-GCaMP3* compound heterozygous BAC transgenic mice, Figure 1A), as described in a previous study(Iwata et al., 2017). When the OB was imaged with two-photon microscopy, we could obtain highly reliable response signals (up to ~200% ΔF/F_0_) for various odorants. Excitatory responses were found in 10-50% of glomeruli in the dorsal OB. Unexpectedly, we often observed reductions in GCaMP signals (up to ~30% ΔF/F_0_ reduction) upon odor stimulation (Figure 1B, Figure S1). This is most likely due to the suppression of spontaneous activity which has also been seen in mitral/tufted (M/T) cell imaging (Economo et al., 2016), thus representing inhibitory responses. The inhibitory responses were shared across fibers within a glomerulus (Figure 1B), suggesting that the inhibition is OR-specific, rather than random. It should be noted that the inhibitory responses would be only seen, in theory, for OSNs with high spontaneous activity. These inhibitory responses have been overlooked in many of the previous studies, most likely due to the low sensitivity and/or slower kinetics of the activity sensors.

**Figure 1.**
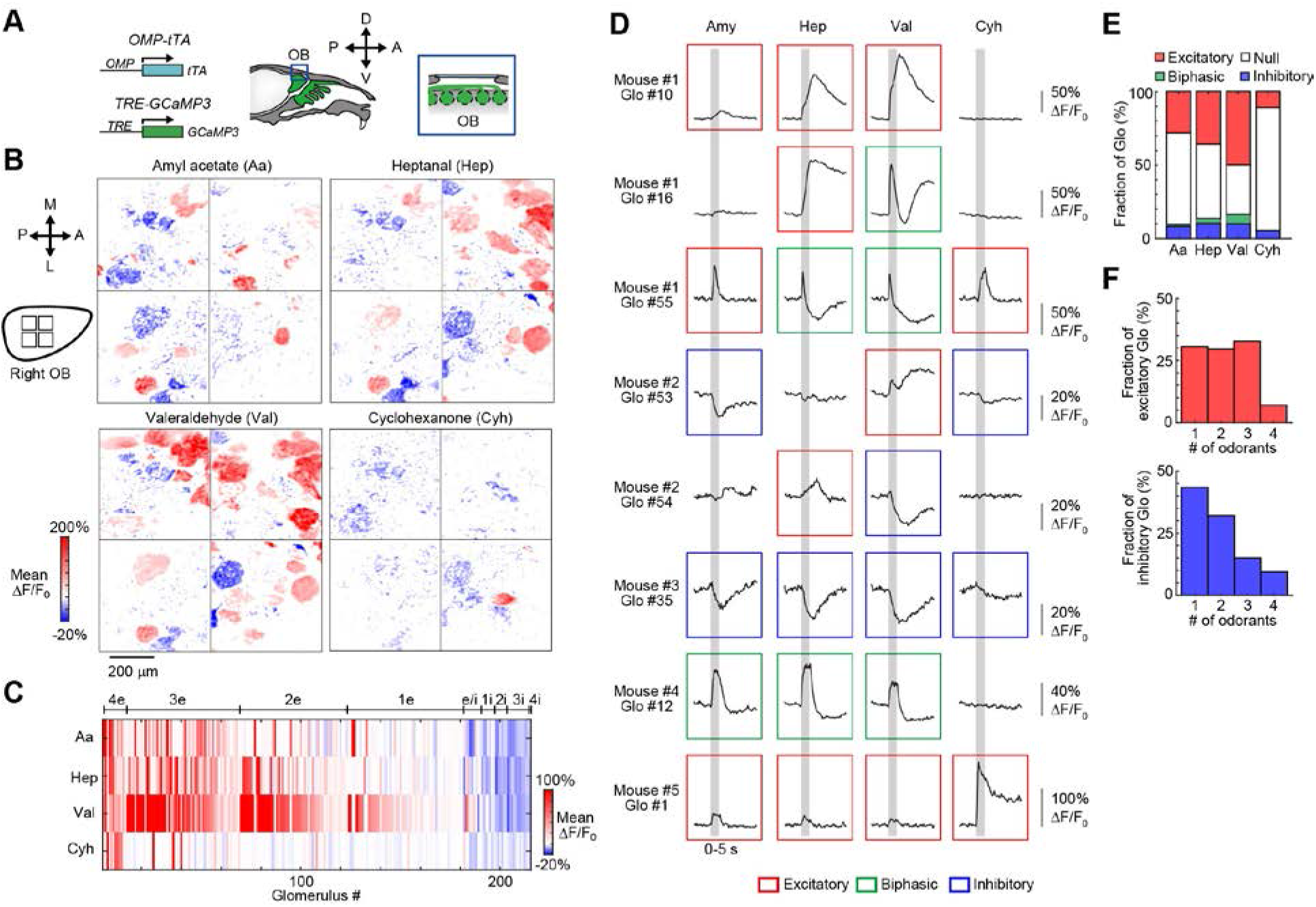
Odor-evoked inhibitory responses in OSN axons. (**A**) Two-photon calcium imaging of OSN axon terminals *in vivo*. OSN-GCaMP3 mice (OMP-tTA BAC Tg crossed with R26-TRE-GCaMP3 BAC Tg) were used for *in vivo* imaging of the OSN axon terminals in the glomerular layer. Only anesthetized mice were analyzed except for Figure S1E, F. Odors were delivered by a custom-made olfactometer (Figure S1H). (**B**) Excitatory (red) and inhibitory (blue) responses at OSN axon terminals in the glomerular layer. Mean ΔF/F_0_ per pixel during the first 10 s from the stimulus onset is shown. Scale bar, 200 μm. A, anterior; P, posterior; M, medial; L, lateral. (**C**) Glomerular and odor specificity of excitatory and inhibitory responses. Both wide and narrow odor tuning was observed for both excitatory and inhibitory responses. Glomeruli showing responses to at least one odorant were analyzed. Glomeruli are clustered based on the number of excitatory (4e-1e) and inhibitory (1i-4i) odors and for bidirectional (e/i) responses. Only significant responses (response mean is >3 SD above/below baseline) were categorized as excitatory or inhibitory. N = 299 glomeruli from 5 mice. 215 glomeruli showing significant responses to at least one odorant are shown. (**D**) Representative glomerular (Glo) responses at OSN axon terminals to odors (Aa, amyl acetate; Hep, heptanal; Val, valeraldehyde; Cyh, cyclohexanone; diluted at 0.5%). Odors were delivered to the nose for 5 s (shown in gray) under freely breathing conditions. Excitatory and inhibitory responses were defined when the response peak and trough were >5 SD above/below baseline, respectively. Biphasic responses demonstrated both excitatory and inhibitory responses. (**E**) Polarity of glomerular responses to each odor. Fractions out of the total number of glomeruli are shown. A small fraction demonstrated biphasic (mostly excitatory-to-inhibitory) responses. (**F**) Tuning specificity of excitatory and inhibitory responses. Data are from 186 and 53 odor-glomerulus pairs for excitatory and inhibitory responses, respectively. Biphasic glomeruli are categorized into both excitatory and inhibitory ones here.

We measured responses to four pure odorants (amyl acetate, heptanal, valeraldehyde, and cyclohexanone, diluted at 0.5%) for 299 glomeruli (5 mice in total) and inhibitory responses were commonly observed for all the four odorants (Figure 1B-E; data for more odorants are described in Figure S1). These responses were reproducible across 3 trials (Figure S1B). Similar to the excitatory responses, the inhibitory responses were broadly tuned in some, but narrowly tuned in other glomeruli (Figure 1F). In this dataset, 60.5% and 8.4% of glomeruli demonstrated excitatory and inhibitory responses to at least one odorant, respectively (mean response is >3 standard deviations (SD) above/below baseline). A substantial fraction of glomeruli (3.0%) demonstrated excitation to some and inhibition to other odorants (e/i fraction in Figure 1C). Some glomeruli demonstrated more complex biphasic responses, showing excitatory-to-inhibitory responses (Figure 1D, E, S1B, D). While most of our experiments were performed under anesthesia, we also observed inhibitory and biphasic responses in OSN axons in awake animals (Figure S1E, F).

It has been known that ~50% of OSNs show mechanosensory responses (Grosmaitre et al., 2007; Iwata et al., 2017). In the *in vivo* situation, sniffing alone produces airflow-dependent mechanical responses in OSNs. Using artificial sniffing, we examined the responses to changes in the nasal airflow speed. Again in this experiment, we observed both excitatory and inhibitory responses to increased airflow in OSN axon terminals (Figure S1G, I-L).

### Inhibitory responses at OSN axon terminals remained without presynaptic inhibition

Next, we investigated the origin of the inhibitory responses seen at OSN axon terminals. The inhibitory responses to odors may be derived from presynaptic lateral inhibition within the OB circuit. Like photoreceptor cells in the retina, OSN axons receive presynaptic inhibition from OB interneurons. A previous study using spH mice and pharmacology indicated that presynaptic inhibition occurs only within a glomerulus, and lateral presynaptic inhibition plays little or no role in odor responses (McGann et al., 2005). However, another study suggested a role for presynaptic lateral inhibition (Fleischmann et al., 2008). A subset of juxtaglomerular interneurons in the OB (GABAergic and dopaminergic short axon cells) are known to innervate multiple glomeruli, and thus may be a potential source for interglomerular lateral inhibition (Kosaka and Kosaka, 2008). We, therefore, examined whether interglomerular presynaptic inhibition could play a role by using conditional mutant mice.

It is known that GABA and dopamine play major roles in the presynaptic inhibition of OSN axon terminals (Aroniadou-Anderjaska et al., 2000; Ennis et al., 2001; McGann et al., 2005). GABA-dependent presynaptic inhibition is mediated by the GABA_B_ receptor. A functional GABA_B_ receptor is a heterodimer consisting of GABA_B1_ and GABA_B2_ subunits (Bettler et al., 2004). We, therefore, generated a conditional mutant mouse line for GABA_B1_, which constitutes a GABA-binding subunit. As for dopamine, the D2 receptor is known to mediate presynaptic inhibition (Ennis et al., 2001). We, therefore, generated a conditional mutant mouse line for D2 receptor (see Figure S2 and Methods for details). Crossed with the *OMP-Cre* knock-in line (Li et al., 2004), we generated OSN-specific double knockout animals for both GABA_B1_ and D2 receptor (Figure 2A; *OMP-Cre*^+/−^; *Gabbr1*^*fl/fl*^; *Drd2*^*fl/fl*^). Immunostaining of OB sections from the conditional mutant mice showed a much-reduced expression of GABA_B_ and D2 receptors (Figure S2H, I). As Cre is highly expressed in all mature OSNs, the weak remaining signals are most likely derived from OB neurons. We examined odor responses using OSN-GCaMP3 (i.e., *OMP-tTA; TRE-GCaMP3; OMP-Cre*^+/−^; *Gabbr1*^*fl/fl*^; *D2R*^*fl/fl*^) mice. We found that the fraction of inhibitory glomeruli is reduced (8.6% in WT and 4.1% in cKO, *p* = 1.4 ×10^−4^, Chi-square test), but not abolished in the mutant mice (Figure 2B).

**Figure 2.**
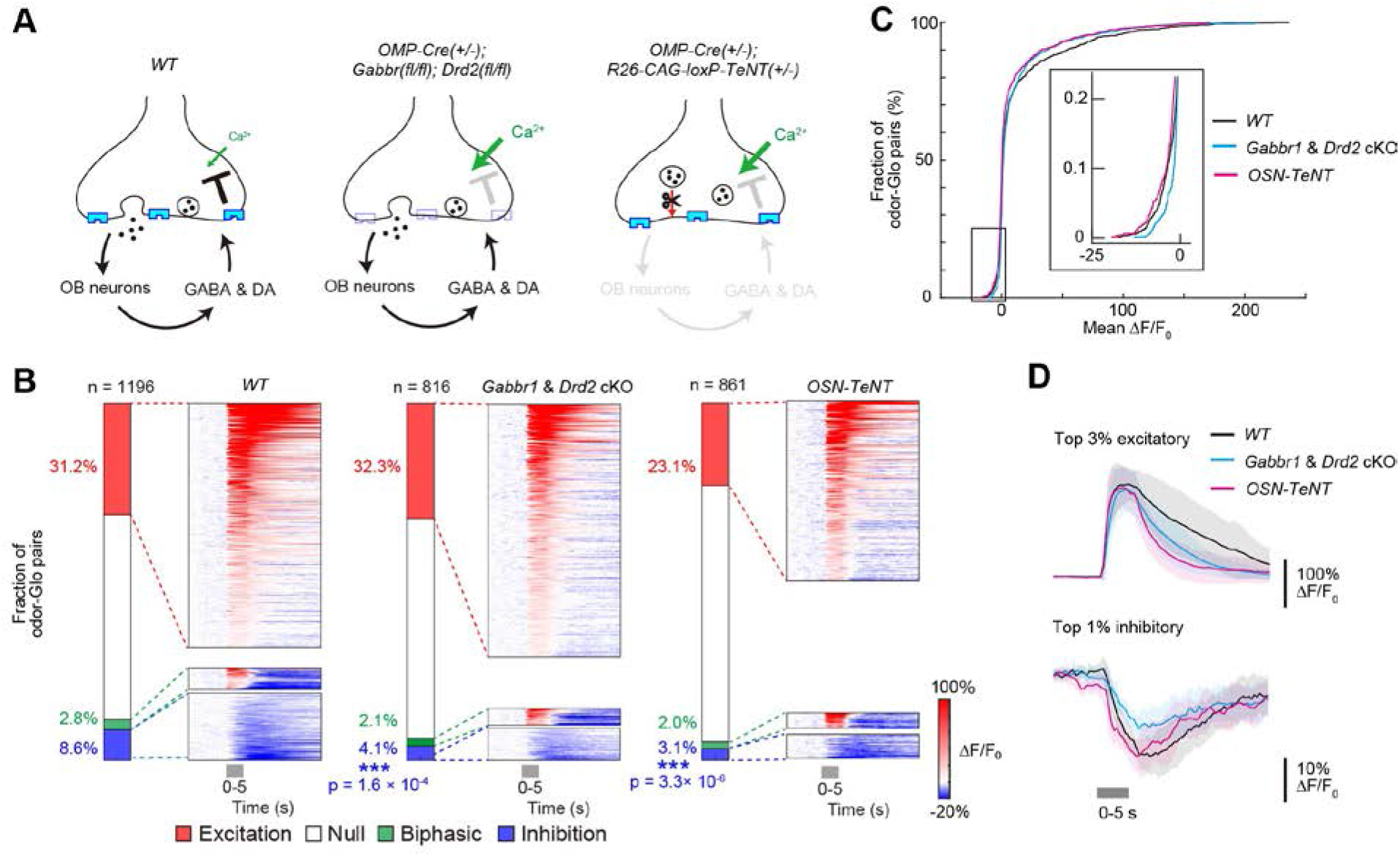
Inhibitory responses at OSN axon terminals without known types of presynaptic inhibition. (**A**) Schematic representation of conditional double knockout and OSN-TeNT mutant mice. We generated OSN-specific *Gabbr1* and *Drd2* mutant mice (*OMP-Cre*^+/−^; *Gabbr1*^*fl/fl*^; *Drd2*^*fl/fl*^) (See Figure S2 and Methods for details). OSN axon terminals of *Gabbr1* and *Drd2* cKO mice cannot receive presynaptic inhibition mediated by GABA and DA. We also analyzed OSN-specific TeNT knock-in mice (OSN-TeNT). OSN axons in OSN-TeNT mice cannot release synaptic vesicles (Figure S3A-D), and thus do not receive presynaptic inhibition from OB interneurons. (**B**) Excitatory and inhibitory glomerular responses (response peak/trough was >5 SD above/below baseline, respectively) in the mutant mice. Amyl acetate, heptanal, valeraldehyde, and cyclohexanone (diluted at 0.5%) were tested. Stacked bar plots indicate the fraction of response types in each mouse line (left). Odor-evoked responses across a population of odor-glomerulus pairs sorted by their response amplitude (right). The *y*-axis represents the odor-glomerulus pairs. WT, N = 1196 odor-glomerulus pairs from 5 mice; *Gabbr1/Drd2* cKO, N = 816 odor-glomerulus pairs from 4 mice; OSN-TeNT, N = 861 odor-glomerulus pairs from 6 mice. p*** < 0.001 compared to WT (Chi-square test). (**C**) Cumulative histogram of response amplitudes. The inset includes expanded *x*- and *y*-axes to display inhibitory response. (**D**) Temporal kinetics of excitatory and inhibitory responses. For a fair comparison across genotypes, the top 3% excitatory and top 1% inhibitory glomeruli (based on the mean response) were used to show the averaged excitatory and inhibitory response kinetics, respectively. WT, N = 34 and 11 odor-glomerulus pairs; *Gabbr1/Drd2* cKO, N = 24 and 8 odor-glomerulus pairs; OSN-TeNT, N = 25 and 8 odor-glomerulus pairs for excitatory and inhibitory responses, respectively. See also Figure S3E-O.

Although GABA and dopamine have been considered to play major roles in presynaptic inhibition in OSN axons, we cannot rule out the contribution of unknown types of presynaptic inhibition mediated by the OB neurons. We, therefore, generated OSN-specific tetanus toxin light chain (TeNT) knock-in mice, in which synaptic transmission from all OSNs is blocked (Figure 2A, *OMP-Cre; R26-CAG-loxP-TeNT*, OSN-TeNT hereafter) (Fujimoto et al., 2019; Sakamoto et al., 2014). These anosmic mice showed poor growth and survival rate during the lactation period but showed almost normal body weight in the adult. Using an M/T cell-specific GCaMP6f mouse line (Dana et al., 2014; Iwata et al., 2017), we confirmed that the odor responses in M/T cells are almost completely shut off in OSN-TeNT mice; instead, slow synchronized activity was found in M/T cells by unknown reasons (Figure S3A-D). Because neurotransmission from OSNs is abolished, all kinds of presynaptic inhibition from OB neurons should be eliminated in this mouse line. Again in OSN-TeNT mice, the fraction of inhibitory glomeruli was reduced compared to wild-type mice (3.1%, *p* = 3.3 × 10^−6^, Chi-square test). The temporal kinetics of excitatory responses were also affected in the mutant mice (Figure 2D, S3E-O). However, a substantial fraction of glomeruli still demonstrated inhibitory responses to odor stimuli in OSN axon terminals (Figure 2B).

**Figure 3.**
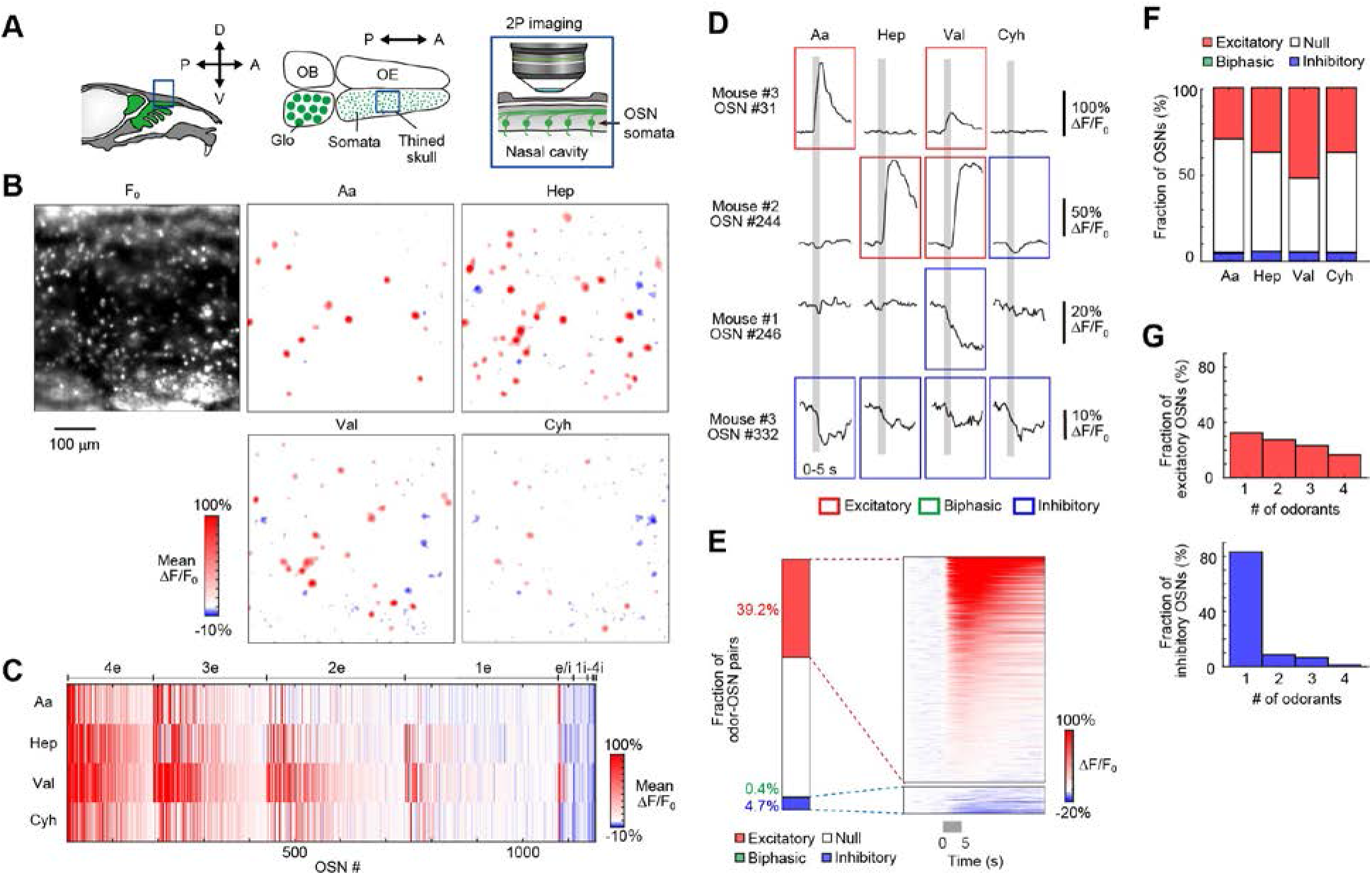
Inhibitory responses at OSN somata in the OE. (**A**) Two-photon calcium imaging of OSN somata in the OE *in vivo*. OSNs in the dorsal OE (zone 1) were imaged through the thinned skull. (**B**) A representative image of basal GCaMP3 fluorescence and widespread inhibitory responses in the OE. Excitatory (red) and inhibitory (blue) responses (response mean) are shown. Scale bar, 100 μm. See also Movie S1. (**C**) Responsive OSNs are clustered based on the number of excitatory (4e-1e) and inhibitory (1i-4i) odors, and for bidirectional (e/i) responses. Only significant responses (response mean was >3 SD above/below baseline) were categorized as excitatory or inhibitory. OSNs showing responses to at least one odorant were analyzed. N = 1,617 OSNs from 4 mice. 1,159 OSNs showing significant responses to at least one odorant are shown. (**D**) Representative response profiles of OSNs to odors. Odors were delivered to the nose for 5s (shown in gray). Traces for both excitatory and inhibitory responses are shown. Excitatory and inhibitory responses were defined for >5 SD changes above/below baseline. Biphasic ones demonstrated both excitatory and inhibitory responses. (**E**) Polarity of odor-evoked responses analyzed for total OSNs. Biphasic responses were not evident in the OE. (**F**) Fractions of OSN somata showing excitatory and inhibitory responses. N = 6,468 odor-OSN pairs. (**G**) Tuning specificity of excitatory and inhibitory responses. Inhibitory responses were more narrowly tuned than in glomeruli. Data are from 1,106 and 137 OSNs for excitatory and inhibitory responses, respectively.

It is possible that homeostatic plasticity has obscured some aspects of presynaptic inhibition under the chronic inactivation present in these mutant mice. Nonetheless, the results so far strongly suggest that a substantial fraction of inhibitory responses is generated within OSNs, without any contribution from OB interneurons.

### Inhibitory responses at OSN somata in the OE

A straightforward way to confirm this possibility is to record odor responses in the OSN somata in the OE *in vivo*. We have recently established *in vivo* two-photon imaging of OSN somata in the OE (Figure 3A) (Iwata et al., 2017). As we used transgenic OMP-tTA to drive TRE-GCaMP3 in our OSN-GCaMP3 mice, only a subset of OSNs (~60%) expresses GCaMP3, allowing for the separation of most of the somatic signals. Due to the high basal fluorescence of GCaMP3 (Figure 3B), we were able to record various types of odor responses (Movie S1). Here we recorded OSN soma responses using the same set of odorants as used for OSN axon imaging (amyl acetate, heptanal, valeraldehyde, and cyclohexanone, diluted at 0.5%). Unlike the vomeronasal organ (Leinders-Zufall et al., 2000; Xu et al., 2016), a single odorant often activated a dense population of OSNs (20-50%) in the OE (response peak is >5 SD above baseline). We also found that 4.7% of OSNs show inhibitory responses at their somata (response troughs >5 SD below baseline) (Figure 3B-F). Biphasic responses were rarely found at the OSN somata. Inhibitory responses at OSN somata were more narrowly tuned than in OSN axon terminals (Figure 3G). However, as the fluorescence signals are much weaker than in the glomeruli, it is difficult to compare them quantitatively (Figure S4A-E).

Similar to the excitatory responses, the inhibitory responses of OSNs were concentration-dependent (Figure 4). More OSNs showed inhibitory responses at higher odor concentrations. Most OSNs demonstrated a tonic increase in the amplitude of excitation and inhibition as the concentration increases (Spearman’s correlation coefficient = 0.46 and −0.18; *p* = 4 × 10^−148^ and 1 × 10^−4^, respectively). We did not find evidence for a concentration-dependent excitatory-to-inhibitory switch of response polarity (Figure 4C). It is therefore unlikely that depolarization-induced inactivation of sodium channels underlies the inhibitory responses in OSNs.

**Figure 4.**
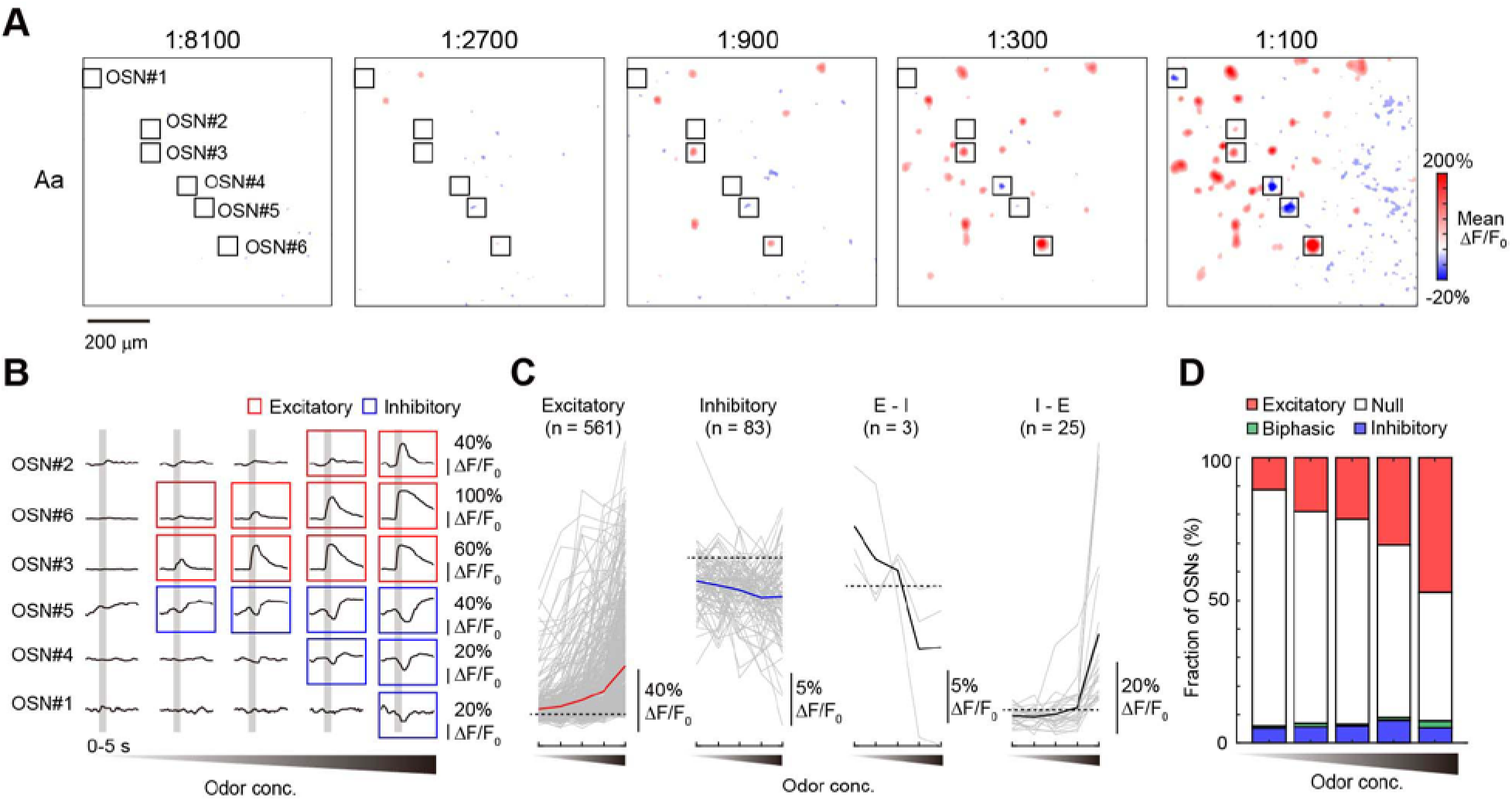
Concentration-dependent inhibition at OSN somata in the OE. (**A**) Concentration-dependent changes in OSN somata responses in the OE *in vivo*. A diluted series of amyl acetate (Aa) (1:8,100, 1:2,700, 1:900, 1:300, and 1:100) were applied. (**B**) Representative response profiles of OSNs to increasing concentrations of odors. (**C**) Response amplitudes for excitatory, inhibitory, excitatory-inhibitory, and inhibitory-excitatory OSNs. Tested odorants are amyl acetate and valeraldehyde (N = 1,118 odor-OSN pairs from 3 mice). Spearman’s correlation coefficient was 0.46 and −0.18 (*p* = 4 × 10^−148^ and 1 × 10^−4^) for excitatory and inhibitory fractions, respectively. (**D**) Fraction of excitatory, inhibitory, and biphasic responses at varying odor concentrations.

### Odorants act as inverse agonists for some ORs in a heterologous assay system

What is the origin of the inhibitory responses seen in OSN somata? In one scenario, the inhibitory responses may be a result of non-synaptic lateral inhibition known as ephaptic coupling. In the *Drosophila* olfactory system, 2-3 OSNs are tightly packed within a sensillum and electrically affect each other (Su et al., 2012; Zhang et al., 2019). This may occur within the tightly packed OE and axon bundles in mice (Bokil et al., 2001); however, each type of OSN is randomly scattered in the OE, suggesting that ephaptic coupling alone cannot account for the OR-specific inhibitory responses found at axon terminals (Figure 1B). We also examined whether inhibitory responses tend to occur when neighboring glomeruli or OSNs show excitatory responses. However, we observed no clear tendency, suggesting that ephaptic coupling plays a minimal role, if any, in the inhibitory responses seen in mice (Figure S4F-H). We, therefore, considered the possibility that the inhibitory responses occur at the OR level, as has also been proposed for *Drosophila* OSNs (Cao et al., 2017; Hallem et al., 2004). Although receptor antagonism in mixture responses has been known (Araneda et al., 2004; Oka et al., 2004; Rospars et al., 2008), inhibitory responses are not fully established for mammalian ORs.

To study inhibitory responses at the receptor level, we used a heterologous assay system. ORs were co-expressed with a chaperone molecule, RTP1S, in HEK293 cells. We examined cAMP responses based on cAMP-response element (CRE) promoter activity using a dual luciferase assay system (See Methods for details) (Saito et al., 2004; Tsuboi et al., 2011). Using the human β2 adrenergic receptor as a control, we confirmed that a known inverse agonist, ICI-118,551, reduces CRE activity in a dose-dependent manner (Figure S5A, B). In the initial screen, we expressed 176 mouse ORs and measured the basal activity without odorants. We then focused on the 17 ORs with the highest basal activity, as it would be easier to find inhibitory responses for these ORs. We then examined the responses of these ORs to 9 odorants. We found that Olfr644 (also known as MOR13-1) and Olfr160 (MOR171-3) show inhibitory responses to odorants. For Olfr644, benzaldehyde acted as an agonist, but heptanal and amyl acetate were inverse agonists showing a reduction in CRE activity (Figure 5A). Heptanal at 300 μM reduced basal CRE activity by ~50%, which was comparable to the inverse agonism activity of ICI-118,551 toward human β2 adrenergic receptor. Similarly, for Olfr160, acetophenone was an agonist, but ethyl hexanoate was an inverse agonist (Figure S5C). More pronounced inhibitory effects were seen when inverse agonists were applied together with agonists (Figure 5B). Thus, at least in part, the inhibitory responses originate from inverse agonism at the receptor level.

**Figure 5.**
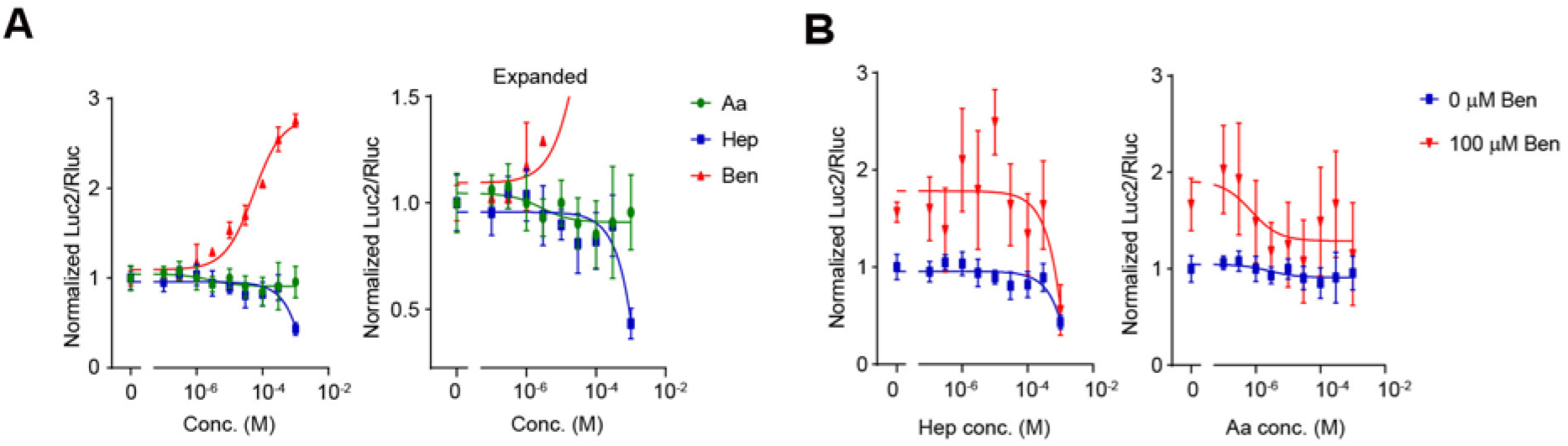
Inhibitory responses in a heterologous assay system for ORs. (**A**) Dose-response curves of Olfr644 to amyl acetate (Aa), heptanal (Hep), and benzaldehyde (Ben). ORs were expressed in HEK293 cells with a chaperone molecule, RTP1S. A dual luciferase assay was used to measure cAMP response element (CRE) promoter activity. Control experiments with the human β2-adrenergic receptor is shown in Figure S5A, B. Benzaldehyde was an agonist for Olf644. However, heptanal and amyl acetate acted as inverse agonists for Olfr644. Similarly, ethyl hexanoate acted as an inverse agonist for Olfr160, whereas acetophenone was an agonist (Figure S5C). (**B**) Inhibition in the presence of agonist. The dose-response curves of two inverse agonists (Hep and Aa) were determined in the presence of 100 μM benzaldehyde.

### Extensive antagonism and synergy in odor mixture responses in OSNs

Natural odors are often composed of multiple odorants. However, they are not always perceived as a simple combination of its components. The logic behind odor mixture perception is an important issue in the field. Previously, it has been widely assumed that odor mixtures are represented as a linear sum of its components at the peripheral level (Fletcher, 2011; Khan et al., 2008; Lin et al., 2006). Therefore, non-linear transformation of odor mixture responses has been considered to occur in the OB (Economo et al., 2016; Yokoi et al., 1995) and in the piriform cortex (Stettler and Axel, 2009) as a consequence of lateral inhibition and/or convergent excitatory inputs.

Here we considered that the inhibitory responses found in OSNs would have an impact on the odor mixture representations at the OSN level. This issue has not been investigated at sufficient detail *in vivo*. We examined the OSN axon responses for amyl acetate (0.5%), valeraldehyde (0.5%), and a mixture of amyl acetate + valeraldehyde (0.5% + 0.5%) (see Figure S1H, S6 and Methods for odor delivery and concentration measurements in mixture experiments). Often we observed that one odorant reduces or abolishes the responses to the other at various degrees (Figure 6). We obtained similar results for various odor mixtures (Figure S7A-C). The suppressive effects of the odor mixture representation were seen not only for glomeruli demonstrating excitatory and inhibitory (E-I) responses to the odor pair but also for excitatory/null (E-N) and even for excitatory/excitatory (E-E) cases (Figure 6C). When we only analyzed glomeruli showing excitatory responses to an odorant (stronger odor) and excitatory or null responses to another (weaker odor; stronger and weaker odors were defined for each glomerulus), 58% of glomeruli demonstrated suppression by weaker odors (Figure 6C, D). These suppressed responses were also seen in OSN-specific GABA_B1_/D2R double knockout mice (data not shown).

**Figure 6.**
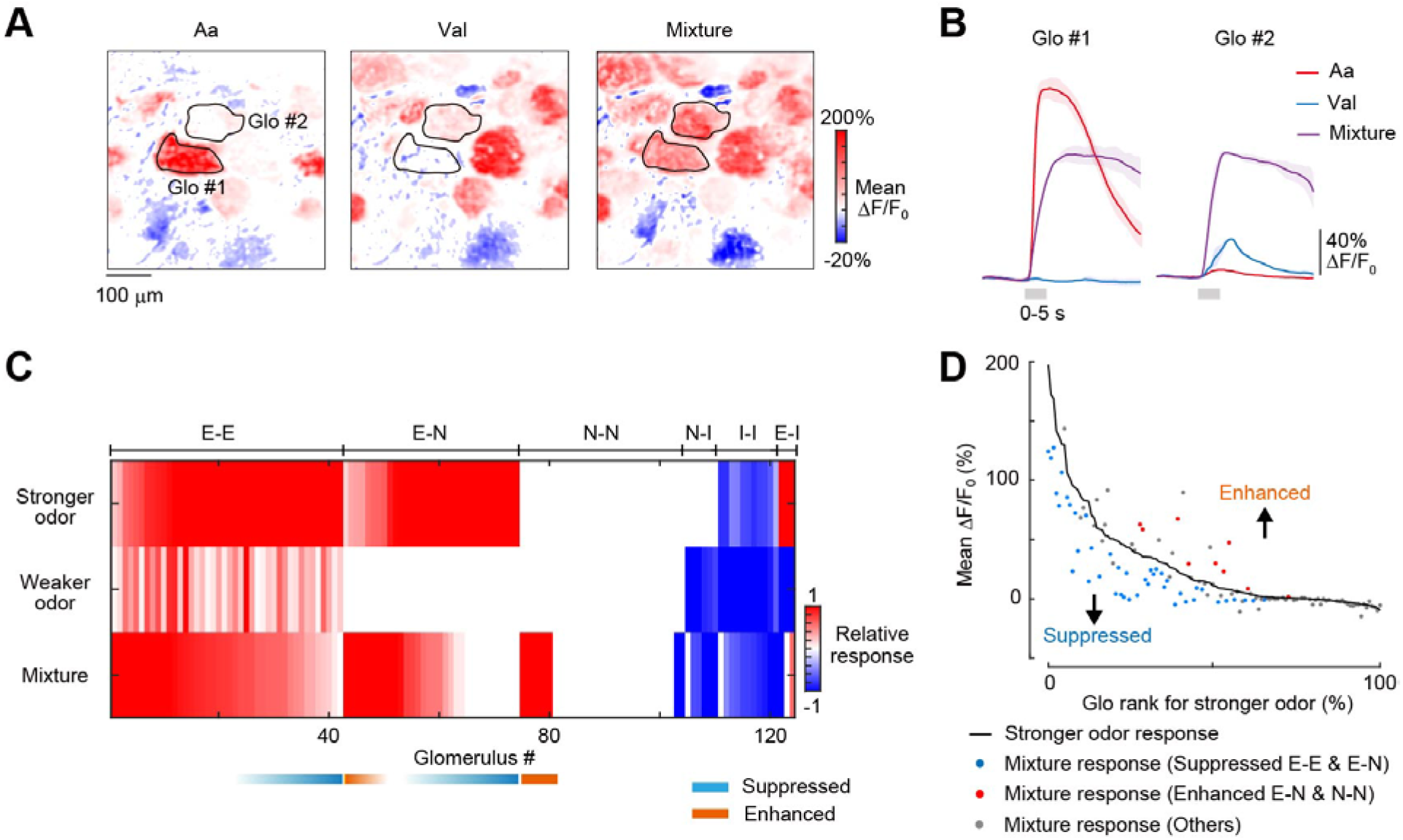
Representation of odor mixtures at OSN axon terminals. (**A**) Responses to amyl acetate (Aa), valeraldehyde (Val), and a mixture of them at OSN axon terminals in the glomerular layer. Mean ΔF/F_0_ per pixel during the first 10 s from the stimulus onset is shown. Scale bar, 100 μm. (**B**) Response profiles of glomerulus #1 and #2 shown in (**A**). Val suppressed the response of glomerulus #1 to Aa. Aa enhanced the response of glomerulus #2 to Val. (**C**) Summary of glomerular responses. A weaker odor was defined as an odor that showed a smaller response in a given glomerulus. A stronger odor was defined as the one showing a larger response. Weaker and stronger odors were defined for each glomerulus. Glomeruli are sorted based on the mean response amplitude for the mixture, and then for the stronger odor. Relative response indicates the normalized response based on the maximum responses among the three odor conditions. Maximum excitatory and inhibitory responses are 1 and −1, respectively. Excitatory (E) and inhibitory (I) responses were defined when response mean >3 SD above baseline. All other glomeruli were defined as null (N) response. Glomeruli were categorized based into E-E, E-N, N-N, N-I, I-I, and E-I types. All glomeruli are shown including ones showing null responses. N = 131 glomeruli from 3 mice. See also raw data in Figure S7A, Aa + Val. (**D**) Glomeruli demonstrating suppressed and enhanced responses in mixture experiments. Glomerulus rank was defined based on the response amplitude to the stronger odor. Two-tailed Student t-test was used to determine whether mixture response was significantly suppressed or enhanced from stronger response in each glomerulus (N = 3 trials each). Suppression was analyzed only for E-E and E-N glomeruli (p < 0.05) and are shown in blue. Enhancement for E-N glomeruli was based on p < 0.05. As for N-N glomeruli, enhancement was defined when the mixture response was judged as excitatory. Enhancement seen for E-N and N-N are shown in red. All other glomeruli are shown in gray.

A more puzzling observation was the supra-linear enhancement of odor responses in the odor mixture experiment, which is known as synergy. For example, the responses to a mixture of amyl acetate + valeraldehyde were much greater than the linear sum of amyl acetate and valeraldehyde responses in some glomeruli (Figure 6A, B). Here we only focused on glomeruli showing excitatory or null responses to an odorant (stronger odor) and null responses to another (weaker odor), to exclude the possible contribution of non-linearity as a result of the GCaMP3 sensor. Nevertheless, 23% of cases demonstrated enhanced responses over the response to the stronger odor (Figure 6C, D).

OSN somata also demonstrated similar suppression and synergy by using odor mixtures (Figure 7, S7D-F). When we only analyzed OSN somata showing excitatory responses to an odorant (stronger odor) and excitatory or null responses to another (weaker odor; stronger and weaker odors were defined for each OSNs), 11% of them demonstrated suppression by the weaker odor (Figure 7C). When we analyzed OSN somata that showed excitatory or null responses to an odorant (stronger odor) and null responses to another (weaker odor), 18% of them demonstrated synergy effect by the weaker odor (Figure 7C, D). Taken together, the results indicate that, the responses to odor mixtures are extensively modulated (either suppressed or enhanced) already in OSN somata.

**Figure 7.**
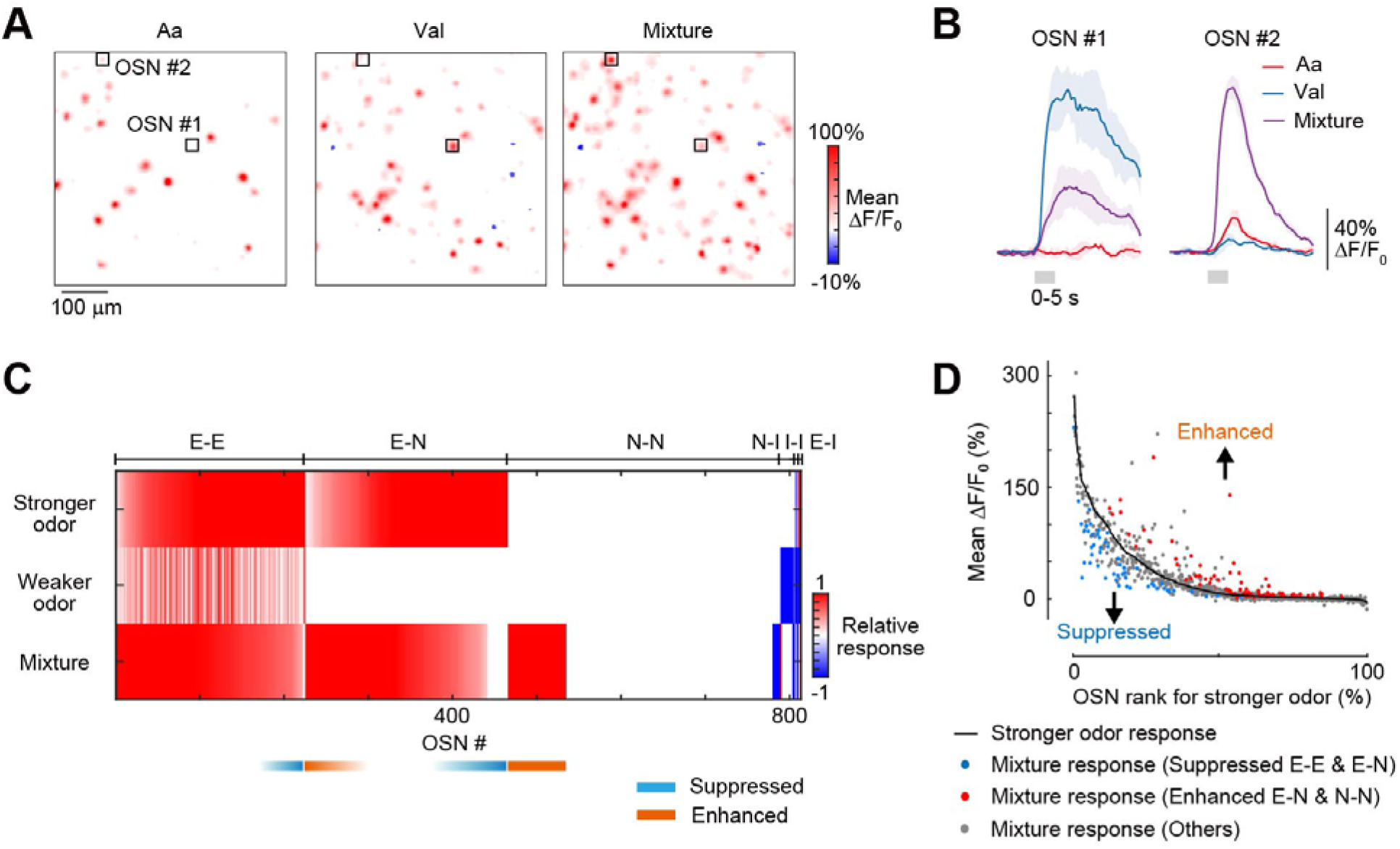
Representation of odor mixtures at OSN somata in the OE. (**A**) Responses to amyl acetate (Aa), valeraldehyde (Val), and a mixture of them at OSN somata in the OE. Mean ΔF/F_0_ per pixel during the first 10 s from the stimulus onset is shown. Scale bar, 100 μm. (**B**) Response profiles of OSN #1 and #2 shown in (A). Aa suppressed the response of OSN #1 to Val. Val enhanced the response of OSN #2 to Aa. (**C**) Summary of OSN responses. A weaker odor was defined as an odor that showed a smaller response in a given OSN. A stronger odor was defined as the one showing a larger response. Weaker and stronger odors were defined for each OSN. OSNs are sorted based on the response amplitude for the mixture, and then for the stronger odor. Relative response indicates the normalized response based on the maximum responses among the three odor conditions. Maximum excitatory and inhibitory responses are 1 and −1, respectively. Excitatory (E) and inhibitory (I) responses were defined when response mean was >3 SD above/below baseline. All other OSNs were defined as null (N) response. OSNs were categorized based into E-E, E-N, N-N, N-I, I-I, and E-I types. All OSNs are shown including ones showing null responses. N = 825 OSNs from 3 mice. See also raw data in Figure S7D, Aa + Val. (**D**) OSNs demonstrating suppressed and enhanced responses in mixture experiments. OSN rank was based on the response amplitude to the stronger odor. Two-tailed Student t-test was used to determine whether mixture response was significantly suppressed or enhanced from stronger response in each glomerulus (N = 3 trials each). Suppression was analyzed only for E-E and E-N OSNs (p < 0.05) and are shown in blue. Enhancement for E-N OSNs was based on p < 0.05. As for N-N OSNs, enhancement was defined when the mixture response was judged as excitatory. Enhancement seen for E-N and N-N are shown in red. All other OSNs are shown in gray.

### Concentration-dependent antagonism vs. synergy biases

What is the logic behind the mixture interactions? We compared mixture responses at high and low odor concentrations of amyl acetate and valeraldehyde (1:30 and 1:300) (Figure 8A). We found that antagonism is predominant when higher concentrations of odors were mixed (suppressed fraction were 15.6% and 0.3% for high and low concentrations, respectively, ; p = 5.8 ×10^−15^, Chi-square test); however, synergy was more prominent when lower concentrations of odors were mixed (enhanced fractions were 2.5% and 28.0% for high and low concentrations, respectively, ; p < 1.0 × 10^−15^, Chi-square test; Figure 8B). This tendency was shared among three animals tested. Thus, the antagonism/synergy bias is concentration dependent.

**Figure 8.**
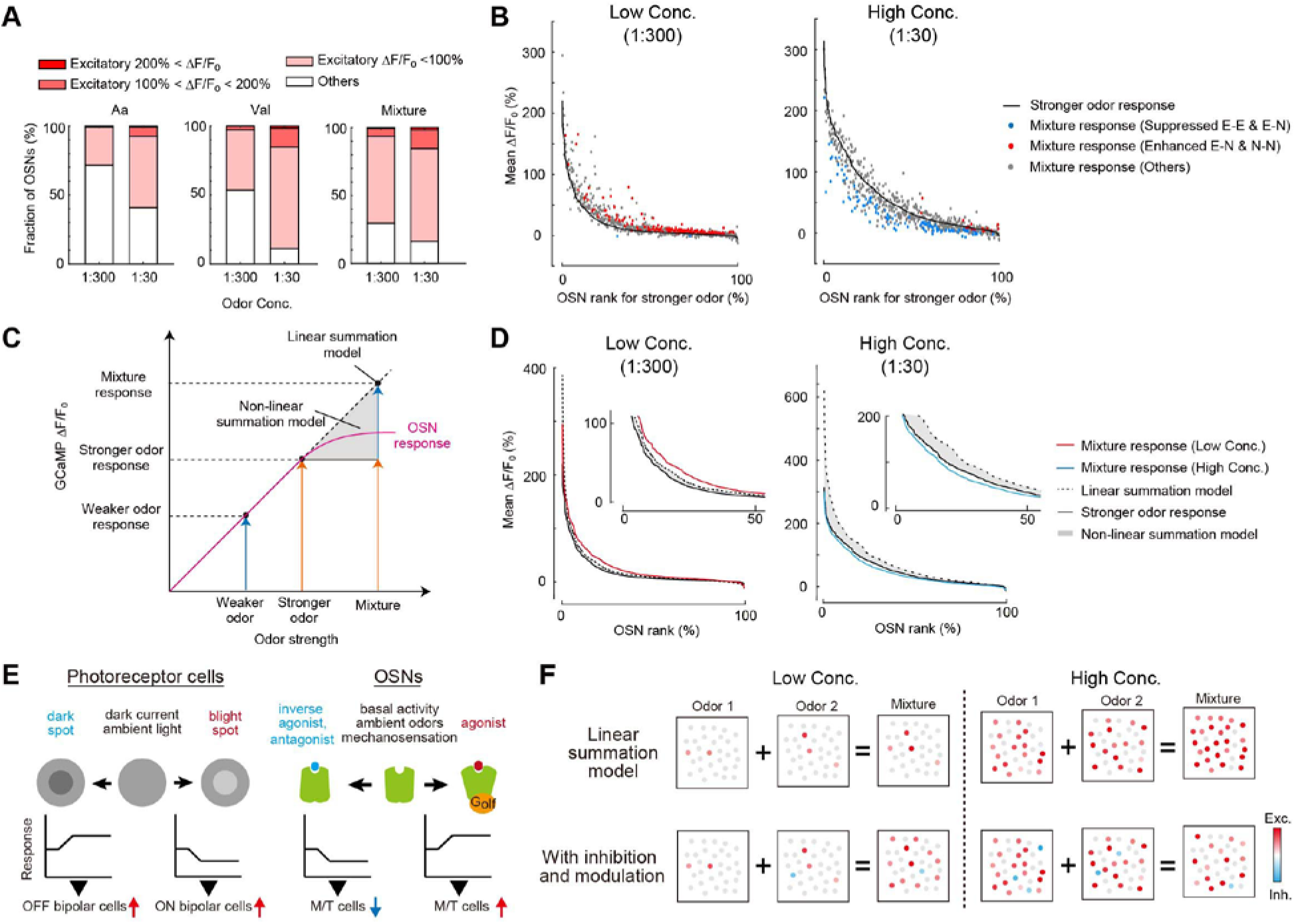
Concentration-dependent odor mixture responses in OSNs. (**A**) Fractions of OSNs showing excitatory responses to high (1:30) and low (1:300) concentrations of amyl acetate (Aa), Valeraldehyde (Val) and their mixtures. N = 680 OSNs from 3 mice. (**B**) Antagonism and synergy found for high and low concentrations of odor mixtures. OSN rank was defined based on the response amplitude to the stronger odor. Suppression seen in E-E and E-N glomeruli are shown in blue, and the enhancement seen for E-N and N-N glomeruli are shown in red. When high concentrations of Aa and Val were mixed (1:30), suppressed responses (antagonism) were often observed: Suppressed fractions were 15.6% and 0.3% for high and low concentrations, respectively (p = 5.8 × 10^−15^, Chi-square test). In contrast, enhanced responses (synergy) were more prominent when odor concentrations were lower (1:300): Enhanced fractions were 2.5% and 28.0% for high and low concentrations, respectively (p < 1.0 × 10^−324^, Chi-square test). (**C**) Linear and non-linear summation models in odor mixture responses. We predicted a range of mixture response profiles based on the responses to each component. The predicted response profiles by the linear and non-linear summation models were compared with the actual responses to the mixtures. (**D**) Responses to odor mixtures compared to linear and non-linear summation models (shown as a gray shade). Predicted and actual responses in OSNs are sorted based on its amplitude. When higher concentrations of odors were mixed, the mixture responses were below the stronger odor curve, indicating that antagonism was dominant in this condition (p = 2.5 × 10^−5^, Wilcoxon rank sum test; p = 7.2×10^−3^, Kolmogorov-Smirnov test). When lower concentrations of odors were mixed, however, the mixture responses were above the linear summation models, which means that synergy was dominant in this condition. (p = 1.6 × 10^−3^, Wilcoxon rank sum test; p = 3.9 × 10^−5^, Kolmogorov-Smirnov test). (**E**) Schematic representation of the bidirectional responses in the visual and olfactory systems. In the visual system, dark current and responses to ambient light set the baseline membrane potential of photoreceptor cells. As a result, photoreceptor cells show bidirectional responses under physiological conditions, which is useful for detecting both bright and dark objects through the ON and OFF pathways. Similarly, in the olfactory system, the basal activity of ORs, ambient odors, and mechanosensation sets the baseline cAMP levels in OSNs. We propose that these bidirectional responses are useful for reliable odor identification in noisy sensory environments. (**F**) The antagonism and synergy bi-directionally control the amplitude and density of odor mixture responses in OSNs, depending on odor concentrations. Natural odors are often composed of multiple components. When low concentrations of odors are mixed, synergistic interactions tend to boost the response amplitude and density. When high concentrations of odors are mixed, however, antagonistic interactions lower the response amplitude and density, avoiding the saturation of the coding capacity. These results were in stark contrast to the classical linear/non-linear summation models. We propose that the widespread inhibition, antagonism, and synergy in OSNs may be useful for robust and efficient coding of odor mixtures at the peripheral stage.

It is technically difficult to examine the functional roles of antagonism and synergy in mixture responses experimentally. Instead, we compared the response amplitude profiles between mixture responses vs. a linear summation model. In the linear summation model, we simply summed responses (ΔF/F) to amyl acetate and valeraldehyde (Figure 8C). However, this should be an overestimate, as the response amplitudes of OSNs may gradually saturate as the receptor inputs increase. We, therefore, also considered a non-linear summation (a monotonically increasing function) model, in which we assumed that the responses to the odor mixtures will be larger than the stronger odor response, but is smaller than the linear summation model (Figure 8C).

When odor concentrations were higher (1:30), the mixture responses were overall smaller and sparser than predicted from the linear summation model, and even below the range predicted from the non-linear summation model (Figure 8D). Thus, the amplitude and density of OSN responses do not increase in odor mixture responses as is predicted from the linear and non-linear summation models, as a result of the widespread antagonism.

In contrast, however, mixture responses were often greater than the linear summation model when odor concentrations were lower (1:300) (Figure 8D). As a result, more OSNs participated in the responses of mixtures than the linear summation model. Similar results were obtained for a different set of odors (Figure S8). Thus, the mixture interactions (antagonism and synergy) bi-directionally control the density and amplitude of mixture responses in OSNs, depending on the odor concentrations. When odor concentrations are higher, antagonism reduces the overall density and amplitudes of the responses; however, synergy tends to boost faint responses when the odor concentrations are lower.

## DISCUSSION

It has long been considered that odorants activate ORs and deliver excitatory inputs to the OB. However, our *in vivo* imaging study demonstrated that odorants elicit not only excitatory but also inhibitory responses in OSNs, contrary to the classical view. These inhibitory responses are widespread phenomena in the mammalian OSNs.

What are the origins of the inhibitory responses in OSNs? In the mammalian retina, horizontal cells are known to mediate lateral inhibition, and this inhibition occurs both presynaptically on photoreceptor cells, and postsynaptically on bipolar cells (Demb and Singer, 2015). We, therefore, considered a possible contribution of presynaptic lateral inhibition by OB interneurons. However, two independent mutant mice lacking known types of presynaptic inhibition still demonstrated inhibitory responses at OSN axon terminals. Moreover, we found that inhibitory responses already exist at the level of OSN somata. These results suggest that a substantial fraction of inhibitory responses originate from the OE. However, as there are some differences between OSN somata and axon terminals (e.g., tuning specificity, fraction of biphasic responses, and the degree of mixture effects), it is likely that presynaptic lateral inhibition also tunes output from OSN axon terminals (Fleischmann et al., 2008).

In the *Drosophila* olfactory system, it has been reported that ephaptic coupling within sensilla and inverse agonism contribute to the inhibitory responses at the OSN somata (Hallem et al., 2004; Su et al., 2012; Zhang et al., 2019). As OSNs and their axons are tightly packed (Bokil et al., 2001), ephaptic coupling could also occur in the mammalian olfactory system, especially when excitatory responses are dense. However, given the random distribution of OSNs in the OE, this does not seem to account for the OR-specific inhibitory responses seen in OSN axon terminals (Figure 1). Moreover, based on the spatial distribution of excitatory and inhibitory responses, we found no evidence for ephaptic coupling in the OE (Figure S4F-H). Using a reconstituted system, we showed that odorants act as inverse agonists at least for some ORs (Figure 5), consistent with earlier observations in isolated OSNs (Rawson et al., 1997; Reisert, 2010). It has been known that the basal activity of ORs is important for OSN development: Many of the ORs have basal activity without odorants and the basal cAMP level contributes to axonal wiring specificity during development (Imai and Sakano, 2008; Imai et al., 2006; Imai et al., 2009; Nakashima et al., 2013). In mature OSNs, the basal activity of ORs is required for generating spontaneous activity in OSNs (Connelly et al., 2015; Reisert, 2010); however, its role in odor coding has been unknown. Our current study suggests that the basal activity of mature OSNs contributes to bidirectional responses, excitatory and inhibitory, to odorants. It is possible that OSNs with high basal activity tend to show inhibitory responses to various odorants, as has been known for M/T cells (Kollo et al., 2014).

However, inverse agonism may be just one mechanism for the inhibition. It is also possible that antagonism has contributed to the apparent inhibitory responses seen in our *in vivo* study. OSNs may respond to ambient odors as well as metabolites in the olfactory mucosa in the physiological condition *in vivo*. Also, the OSNs respond to mechanical stimuli produced by the nasal airflow. Although the exact mechanism is yet to be established, an *in vitro* study suggested that ORs may directly recognize mechanical stimuli, similar to odor recognition (Connelly et al., 2015). Therefore, antagonistic odorants can suppress these responses, as shown in our odor mixture experiments (Figure 6-8). In the visual system, the opsin proteins themselves have extremely low basal activity; however, ambient light allows for bidirectional responses (inhibition and excitation) in photoreceptor cells that are conveyed to ON/OFF pathways, respectively (Figure 8E). Similarly, hair cells in the auditory and vestibular systems are known to show bidirectional responses. The inhibitory responses of OSNs may be useful for contrast enhancement and aid robust odor identification under background odors (Figure 8F).

In the OB, all neurons show excitatory responses towards glutamatergic OSN inputs. However, it has been well known that M/T cell responses are highly dynamic, particularly in awake conditions (Cury and Uchida, 2010; Shusterman et al., 2011). M/T cells show both excitatory and inhibitory responses, contributing to timing-based representation of odor information (Economo et al., 2016; Fukunaga et al., 2014; Iwata et al., 2017; Kollo et al., 2014). Therefore, both excitatory and inhibitory responses in OSNs should contribute to the timing-based odor coding in the OB.

Previously, it has been considered that odor mixtures are represented as a linear sum of its components at the peripheral stage (Fletcher, 2011; Khan et al., 2008; Lin et al., 2006). Antagonism for ORs has been known for a few ORs, including rat I7 and MOR-EG (Araneda et al., 2004; Oka et al., 2004). However, its generality remained elusive, and it had yet to be fully established how OR antagonism impacts olfaction *in vivo*. In the current study, we showed that the responses to odor mixtures are already extensively modulated at the most peripheral level *in vivo*. Responses to an odorant are inhibited by another in ~60% of glomeruli and ~10% of OSN somata, supporting the idea that the extensive antagonism tunes the representation of odor mixtures in OSNs (Figure 7). Our *in vivo* observations are in good agreement with recent studies (including bioRxiv preprints) using reconstituted systems (Pfister et al., 2019), *ex vivo* imaging (Xu et al., 2020), and *in vivo* (de March et al., 2020; Zak et al., 2019). The discrepancy from earlier studies may be due to the sensitivity of the imaging methods, choice of odorants, and their concentrations (dense responses tested in this study vs. only sparse responses tested in earlier studies). Remarkably, our study showed that a single odorant activates up to ~90% of OSNs in the OE *in vivo*, indicating dense coding of these odorants at this stage. Therefore, the odor mixture would easily saturate OSN responses in a linear summation model. The widespread inhibition and antagonism found in this study should be useful to normalize the population responses and avoid saturation for efficient coding of odor mixtures (Reddy et al., 2018) (Figure 8F).

Unexpectedly, a substantial fraction of OSNs (10-20% both in glomeruli and OSN somata) demonstrated supra-linear enhancement in odor responses (synergy) when mixed (Figure 6, 7). This tendency was particularly prominent when lower concentrations of odors were mixed (Figure 8): The synergistic effect contributes to boost otherwise faint population odor responses. Although enhanced responses at axons may be partly due to the disinhibition of OB inhibitory circuits, we could still see the enhanced responses at OSN somata. This may be due to the allosteric effect of an odorant to another, as has been known to occur for various GPCRs including taste receptors (Schwartz and Holst, 2007). A theoretical framework for synergy has been known (Reddy et al., 2018; Singh et al., 2019), but this issue needs to be further investigated experimentally in the future. Potentially, this may explain how a small amount of additional components produces unexpected and huge perceptual effects in fragrances. The bi-directional nature of mixture effects (antagonism vs. synergy) is useful to optimize the OSN response amplitudes and density to the optimum range, which potentially help optimize coding capacity and robustness of the system (Figure 8F).

Together, our results revise a long-standing view in which odor responses are only excitatory and odor mixtures are represented as the linear sum of individual ones in OSNs. Previous studies proposed that the non-linear representation of odor mixtures is the hallmark of the OB and olfactory cortex, but this is, in fact, a widespread phenomenon already present in OSNs. We propose that extensive inhibition, antagonism, and synergy at the peripheral level contributes to the robust odor coding in the mammalian olfactory system.

## EXPERIMENTAL MODEL AND SUBJECT DETAILS

### Mouse strains

All animal experiments were approved by the Institutional Animal Care and Use Committee (IACUC) of the RIKEN Kobe Branch and Kyushu University. BAC transgenic *OMP-tTA* (line #3; Accession No. CDB0506T: http://www2.clst.riken.jp/arg/TG%20mutant%20mice%20list.html; MGI:5922006) crossed to BAC transgenic *TRE-GCaMP3* mice (high copy line; Accession No. CDB0505T; MGI:5922007) were described in a previous study (Iwata et al., 2017). *R26-CAG-LoxP-TeNT* knock-in (RBRC05154; RRID:IMSR_RBRC05154) (Sakamoto et al., 2014), *OMP-Cre* knock-in (JAX #006668; RRID:IMSR_JAX:006668) (Li et al., 2004), and *Thy1-GCaMP6f* Tg (line GP5.11) (JAX #024339; RRID:IMSR_JAX:024339) (Dana et al., 2014) have been described previously. Cre is expressed in all mature OSNs in OMP-Cre mice (Li et al., 2004). *Drd2*^*tm1a*^ and *Gabbr1*^*tm1a*^ mice were produced from ES clones obtained from EUCOMM and KOMP, respectively. Among the three clones each for *Drd2*^*tm1a*^ and *Gabbr1*^*tm1a*^, only *Drd2*^*tm1a(EUCOMM)Hmgu*^ (HEPD0654_5_D11) and *Gabbr1*^*tm1a(KOMP)Wtsi*^ (EPD0730_1_G04) were transmitted to the germline. Southern blotting was used to confirm correct gene targeting. These mice were then crossed to *Flp* mice (JAX# 003800; RRID; IMSR_JAX: 003800) to delete *lacZ-Neo*^*r*^ cassette and obtained *Drd2*^*tm1c*^ and *Gabbr1*^*tm1c*^ (Figure S2). Genotyping primers were 5’-TCACCCTCCAGCCTGCCTAC-3’ and 5’-CGTCGCGATGTGAGAGGAGA-3’ for *OMP-tTA* mice; 5’-AACCGTCAGATCGCCTGGAG-3’ and 5’-CGGTACCGCCCTTGTACAGC-3’ for *TRE-GCaMP3* mice; 5’-TGTGGAAGGCAATTCTGAGAGG-3’ and 5’-CCACTTTGTACAAGAAAGCTGGGTCT-3’ for *Gabbr1*^*tm1c*^ allele; 5’- GTTGCCTTCCCCCTCTTGCT-3’ and 5’-CCACTTTGTACAAGAAAGCTGGGTCT-3’ for *Drd2*^*tm1c*^ allele (the primer sites are indicated in Figure S2). *OMP-Cre* was in a 129/C57BL/6 mixed background and all other lines were in a C57BL/6N (RRID:MGI:5295404) background. Both male and female mice were used.

## METHOD DETAILS

### Southern blotting

Genomic DNA (10 μg) extracted from the mouse tail was digested with restriction enzymes: *KpnI* (Toyobo), *ApaI* (NEB, R0114S), *SpeI* (NEB, R0133S), and *XhoI* (NEB, R0146S). Electrophoresed agarose gels were treated with 0.4% (v/v) HCl and then alkaline transferred with 0.4 M NaOH to Biodyne Plus Membrane (0.45 μm, Pall Corporation, #60406). Hybridization with DIG-labelled DNA probes was performed in Church’s buffer at 65°C overnight. Blocking was performed with 1.5% Blocking Reagent (Roche, #11096176001), and 0.1% Anti-Digoxigenin-AP Fab fragments (Roche, #11093274910) in blocking buffer was used to detect DIG. Chemiluminescence reaction was performed with CDP-Star substrate (Thermo Fisher Scientific, #11685627001), and detected with an image reader (Fujifilm, LAS-3000 mini). DNA Probes were labeled with DIG-High prime (Roche, #1585606). The locations of DNA probes are indicated in Figure S2. Primer sequences for the probes are 5’-GGAAACTGGGAGGTGGCTCA-3’ and 5’-ATCAGGCTTGGCTTGGCTTG-3’ for *Drd2* upstream; 5’ -CTGGGGAGACCACCAGCAGT-3’ and 5’ -CATGGATCCAACCCCAGAGC-3’ for *Drd2* downstream; 5’ -GTCAGTTCTTGGCCGCAAGC-3’ and 5’-ACTTTCCGGGCTTCGGTCTC-3’ for *Gabbr1* upstream; 5’-TCCTGCAGTTCCATCCACCA-3’ and 5’ -CCACCCGAGTTTTGGGATTG-3’ for *Gabbr1* downstream; 5’ -GCGATACCGTAAAGCACGAG-3’ and 5’ -GCTTGGGTGGAGAGGCTATT -3’ for *Neo*^*r*^ probe.

### Histochemistry

Mice were deeply anesthetized with an overdosing i.p. injection of pentobarbital. After intracardiac perfusion with 4% PFA (Nacalai Tesque, #26126-25) in PBS, the OB was dissected and post-fixed in 4% PFA in PBS overnight. The OB was then cryoprotected with 30% sucrose and then embedded in OCT Compound (Sakura, #4583). Frozen sections were cut at 18 μm thick with a cryostat (Leica). Antigen retrieval was performed by heating in a microwave oven for 5 min in Histofine antigen retrieval solution pH9 (Nichirei, #415201). Sections were pretreated with 4% PFA in PBS and 5% Donkey serum (Sigma, #D9663) in PBS with 0.1% Triton-X100. Rabbit anti-Drd2 (Millipore, AB5084P, RRID: AB_2094980) and guinea pig anti-Gabbr1 (Millipore, AB2256, RRID: AB_11210385) were used at 1:100 and 1:500 dilutions, respectively. AlexaFluor647-conjugated secondary antibodies (Thermo Fisher Scientific, A31573 and A21450, RRID: AB_2536183 and AB_141882) were used at 1:200 dilutions. Sections were counterstained with DAPI (Dojindo, #D523). Immunofluorescence was imaged with an inverted confocal laser-scanning microscope (Olympus, FV1000) using a 20x dry objective lens (Olympus).

### *In vivo* two-photon imaging

*In vivo* imaging of the OB and OE was performed as described previously (Iwata et al., 2017). Adult mice (8-16 weeks of age) were used for imaging. Surgery and imaging under anesthesia was performed under ketamine (Daiichi-Sankyo) – xylazine (Bayer) (80 mg/kg and 16 mg/kg, respectively) anesthesia. During surgery and imaging, the depth of anesthesia was assessed by the toe-pinch reflexes, and supplemental doses were added when necessary. For the imaging of the OB, a craniotomy (2-3 mm in diameter) was made over the dorsal OB leaving the dura mater intact. The OB was covered with a thin layer of silicone sealant (Kwik-Sil, WPI, #KWIK-SIL) and a 3 mm diameter circular coverslip (Matsunami, custom-made), which was secured with super-glue and dental cement (Shofu). For the imaging of the OE, the dorsal part of the D zone (zone 1) OE was imaged through the thinned skull. We used a dental drill with a 1 mm drill tip to evenly thin the skull above the OE. PBS was applied to the thinned area during the imaging. For head-fixation, a custom aluminum head bar (4 × 22 mm) was glued to the skull behind the cranial window. Body temperature was maintained with a heating pad (Akizuki, M-08908). In some experiments (Figure S1E, F), we imaged awake mice as described previously (Guo et al., 2014; Iwata et al., 2017). Water-restriction was started 2-3 days after surgery. Mice under water-restriction were acclimated to head-fixation in an acrylic tube within 3-4 days, 30 min each. Mice were kept in the acrylic tube during the imaging. Imaging with artificial sniffing was performed as described previously (Iwata et al., 2017). The silicon tube inserted into the trachea was connected to a solenoid valve, a flow meter, and an air suction pump in all the experiments. The solenoid valves for nasal airflow and odor delivery were regulated through relay circuits and the computer programs were written in LabVIEW (National Instruments, RRID:SCR_014325). We used a two-photon microscope, model FV1000MPE (Olympus) with Fluoview FV10-ASW software (Olympus, RRID:SCR_014215) and a 25x objective lens (Olympus, XLPLN25XWMP). GCaMP3 was excited at 920 nm with Insight DS Dual (Spectra-Physics).

### Olfactometry

Olfactory stimulation using an olfactometer was described previously (Iwata et al., 2017). The olfactometer consists of an air pump (AS ONE, #1-7482-11), activated charcoal filter (Advantec, TCC-A1-S0C0 and 1TS-B), and flowmeters (Kofloc, RK-1250) (Figure S1H). Odorants were diluted at a defined concentration (described in Figure legends) in 1 mL mineral oil in a 50 mL centrifuge tube. In the odor mixture experiments, two odorants were diluted in the mineral oil. Odor concentrations in saturated vapor were measured by a semiconductor-based odor sensor, OMX-SRM (AS ONE, #2-3484-01), equipped with two odor detectors (detector X and Y). This sensor returns the Euclidean distance between the measured point (X value, Y value) and the origin (0,0), and angle of the line in the XY coordinate. Since the Euclidean distance calculated by detector X and Y is nonlinear against odor concentration, we used the value of detector X to calculate the relative odor concentration. We confirmed that the odor concentrations in the saturated vapor were linearly correlated with the concentrations in the mineral oil (Figure S6). Saturated odor vapor in the centrifuge tube was delivered to a mouse nose with a Teflon tube. The tip of the Teflon tube was located 1 cm from the nose of the animals. Diluted odors were delivered at 1 L/min. Rapid onset and offset odor delivery were confirmed in a previous study (Iwata et al., 2017). Intervals between trials were 2 min and 5 min for lower (< 1%) and higher concentrations (> 1%), respectively, to avoid adaptation. Odorants were purchased from FUJIFILM-Wako (amyl acetate, #018-03623) and Tokyo Chemical Industry (heptanal, #H0025; valeraldehyde, #V0001; cycrohexanone, #C0489), stored at 4°C and diluted in mineral oil just prior to use. 50 mL centrifuge tubes and Teflon tubes were replaced with a new one every time we change odors.

### Luciferase assay

The luciferase assay with HEK 293 cells (293AAV cell, Cell Biolabs) was performed as described previously (Tsuboi et al., 2011). ORs and *Rtp1* (coding for RTP1S) genes were PCR amplified from cDNA prepared from C57BL6/N mouse OE and subcloned into pME18S-F-R vector and pcDNA3, respectively. In the pME18S-F-R vector, OR genes were expressed under the SRα promoter. The OR has a N-terminal FLAG-tag and 20 aa rhodopsin tag to facilitate cell surface expression. Cells were grown in DMEM supplemented with 10% FBS and 1% Penicillin/Streptomycin. Cells seeded in 96-well white-well plates (60% confluent) were transfected with pME18S-F-R-OR (125 ng/well), CRE-Luc2P (25 ng/well; Promega pGL4.29; #E8471), TK-hRluc (25 ng/well, Promega pGL4.74; #6921) and pcDNA3-RTP1s (25 ng/well) using PEI Max (Polysciences, Inc., #24765-1). Twenty-four hours after transfection, the medium was replaced with DMEM with odor ligands and without serum. To avoid the contamination of odors to other wells, the wells were separated by at least 3 wells from the ones containing a different stimulation medium and sealed with a plastic film, SealPlate (Excel Scientific, #STRSEALPLT), during incubation. Cells were incubated for 4 hrs, and then Luc2P and hRluc activities were quantified with the Dual-Glo Luciferase assay system (Promega, #E2920) and a luminometer, model TriStar LB941 (Berthold). Data were collected with Mikro Win 2000 software (Berthold). Data are mean ± SD based on 3 and 6 replicates for agonist and inverse-agonist in one representative experiment, respectively. Hill curve fitting was applied to the dose-response data by using Prism software (GraphPad Software, RRID:SCR_002798). Reproducibility was confirmed by multiple experiments. Newly generated pME18S-F-R-OR plasmids will be deposited to Addgene. The human β2 adrenergic receptor (hβ2AR) was used as a control. To identify inverse agonists for ORs, we have analyzed 176 ORs (Olfr9, Olfr13, Olfr16, Olfr19, Olfr24, Olfr30, Olfr39, Olfr43, Olfr49, Olfr54, Olfr57, Olfr60, Olfr62, Olfr73, Olfr92, Olfr96, Olfr101, Olfr108, Olfr110, Olfr121, Olfr134, Olfr143, Olfr146, Olfr147, Olfr149, Olfr150, Olfr152, Olfr160, Olfr161, Olfr165, Olfr166, Olfr168, Olfr175, Olfr202, Olfr247, Olfr279, Olfr291, Olfr295, Olfr313, Olfr315, Olfr323, Olfr328, Olfr340, Olfr356, Olfr360, Olfr362, Olfr365, Olfr367, Olfr368, Olfr381, Olfr382, Olfr398, Olfr399, Olfr406, Olfr411, Olfr414, Olfr429, Olfr433, Olfr449, Olfr450, Olfr459, Olfr463, Olfr477, Olfr502, Olfr513, Olfr520, Olfr521, Olfr525, Olfr527, Olfr530, Olfr571, Olfr584, Olfr644, Olfr694, Olfr716, Olfr726, Olfr733, Olfr734, Olfr745, Olfr746, Olfr748, Olfr749, Olfr776, Olfr791, Olfr792, Olfr801, Olfr807, Olfr815, Olfr820, Olfr821, Olfr822, Olfr828, Olfr829, Olfr837, Olfr845, Olfr849, Olfr851, Olfr853, Olfr855, Olfr862, Olfr870, Olfr873, Olfr878, Olfr895, Olfr902, Olfr916, Olfr958, Olfr959, Olfr965, Olfr969, Olfr972, Olfr979, Olfr983, Olfr985, Olfr992, Olfr1010, Olfr1019, Olfr1023, Olfr1031, Olfr1032, Olfr1034, Olfr1036, Olfr1046, Olfr1058, Olfr1062, Olfr1079, Olfr1080, Olfr1089, Olfr1100, Olfr1104, Olfr1107, Olfr1120, Olfr1158, Olfr1168, Olfr1184, Olfr1189, Olfr1021, Olfr1085, Olfr1132, Olfr1134, Olfr1184, Olfr1223, Olfr1231, Olfr1233, Olfr1238, Olfr1246, Olfr1253, Olfr1258, Olfr1261, Olfr1264, Olfr1268, Olfr1269, Olfr1276, Olfr1280, Olfr1317, Olfr1323, Olfr1328, Olfr1339, Olfr1344, Olfr1352, Olfr1356, Olfr1395, Olfr1396, Olfr1402, Olfr1411, Olfr1417, Olfr1428, Olfr1440, Olfr1448, Olfr1471, Olfr1477, Olfr1496, Olfr1497, Olfr1502, Olfr1509, Olfr1511). In the initial screen, we identified 17 ORs (Olfr101, Olfr110, Olfr160, Olfr168, Olfr368, Olfr433, Olfr449, Olfr644, Olfr734, Olfr749, Olfr801, Olfr979, Olfr1168, Olfr1328, Olfr1352, Olfr1395, Olfr1411) as showing the highest basal activity without odorants. We then analyzed responses to 9 odorants at 300 μM: amyl acetate (FUJIFILM-Wako, #018-03623), acetophenone (FUJIFILM-Wako, #014-00423), benzaldehyde (Nacalai-Tesque, #04006-62), cyclohexanone (Tokyo Chemical Industry, #C0489), ethyl hexanoate (Tokyo Chemical Industry, #H0108), heptanal (Tokyo Chemical Industry, #H0025), hexanoic acid (Tokyo Chemical Industry, #H0105), valeraldehyde (Tokyo Chemical Industry, #V0001), and methyl valerate (Tokyo Chemical Industry, #V0005). Agonists and inverse agonists were further analyzed for dose-response curves.

### Image data analysis

All image data analysis was performed in MATLAB (Mathworks, RRID:SCR_001622). Lateral drift in time-lapse imaging data was corrected using custom code based on the correlation coefficient. ROIs for glomeruli and somata were manually determined. The ΔF was normalized to the mean intensity for 10 s before stimulus onset (F_0_), and the response amplitude was defined as the peak/trough ΔF/F_0_ during the first 20 s after stimulus onset. The average response map was made based on the mean ΔF/F_0_ during the 10 s after stimulus onset. When the peak or trough ΔF/F_0_ was higher or lower than 5 SD of basal fluctuation level before stimulation onset, the response was categorized as excitatory or inhibitory, respectively. ROIs showing both excitatory peaks and inhibitory troughs were categorized as biphasic. Some glomeruli or OSNs demonstrated tonic increase or decrease of GCaMP fluorescence before odor stimulation. We, therefore, excluded data when a response slope (0-3 s after the stimulus onset) of a glomerulus/OSN was within 50-150% of that before stimulus onset. Temporal median filtering with 3 s window size was applied to the data of OSN somata to make its noise fluctuation level similar to those of glomeruli (Figure S4A, B). To determine the antagonistic effects in odor mixture experiments, we performed two-tailed t-test with stronger odor (one of the two odorants, evoking higher response than the other) and the mixture. We defined weaker odor (showing smaller response) and stronger odor (showing larger response) for each glomerulus. As for the synergy effects in mixture experiments, we compared the linear summation of stronger and weaker odor responses versus the mixture response. Codes are available upon request.

### Statistical analysis

The MATLAB Statistics Toolbox was used for statistical analysis. The number of glomeruli, cells, and mice was described within figure legends. No blinding was performed in data analysis. Chi-squared test was used in Figure 2B. One-way ANOVA and Tukey-kramer post-hoc test was used in Figure S3H-O. Two-tailed Student t-test was used in Figure 6D, 7D, 8B, S7C, S7F, S8B. Wilcoxon rank sum test was used in Figure 8D, S4G, S4H. Kolmogorov-Smirnov test was used in Figure 8D, S8C. Statistical significance was set at P < 0.05. We excluded animals with poor imaging quality but did not perform data exclusion for other reasons.

## DATA AND SOFTWARE AVAILABILITY

Newly generated plasmids, pcDNA3-RTP1s (Addgene #149364), pME18S-F-R-hβ2AR (Addgene #149367), pME18S-F-R-Olfr644 (Addgene #149365), and pME18S-F-R-Olfr160 (Addgene #149366) are available from Addgene. Raw imaging data will be deposited to SSBD:repository (http://ssbd.qbic.riken.jp/).

### Code Availability

All custom-written code is available upon reasonable request by the Lead Contact, Takeshi Imai.

## ACKNOWLEDGMENTS

We thank KOMP and EUCOMM for ES clones (conditional *Drd2* and *Gabbr1*); M. Yokoi (*Pcdh21-Cre*), I. Imayoshi (*R26-CAG-LoxP-TeNT*), K. Svoboda (*Thy1-GCaMP6f*), and P. Mombaerts (*OMP-Cre* knock-in) for mouse strains; Marcus Leiwe for comments on the manuscript. Animal experiments including generation of chimeric mice were supported by the Laboratory for Animal Resources and Genetic Engineering at the RIKEN Center for Life Science Technologies. We appreciate the technical assistance by Mariko Nishihara, Tomoko Ohmine, and from The Research Support Center, Research Center for Human Disease Modeling, Kyushu University Graduate School of Medical Sciences. This work was supported by grants from the PRESTO program of the Japan Science and Technology Agency (JST) (T.I.), the JSPS KAKENHI (23680038, 15H05572, 15K14336, 16K14568, 16H06456, and 17H06261 to T.I., 15K18353 to R.I.), intramural grant from RIKEN Center for Developmental Biology (T.I.), and Grant-in-Aid for JSPS Research Fellow (15J08987 to R.I., and 18J00899 to S.I.).

## AUTHOR CONTRIBUTIONS

These authors contributed equally: Shigenori Inagaki, Ryo Iwata S.I. performed experiments and analyzed data. R.I. generated knockout and transgenic mice and performed initial rounds of imaging experiments. M.I. assisted the OR assay. T.I. supervised the project. S.I. and T.I. wrote the manuscript.

---

**Figure S1.**
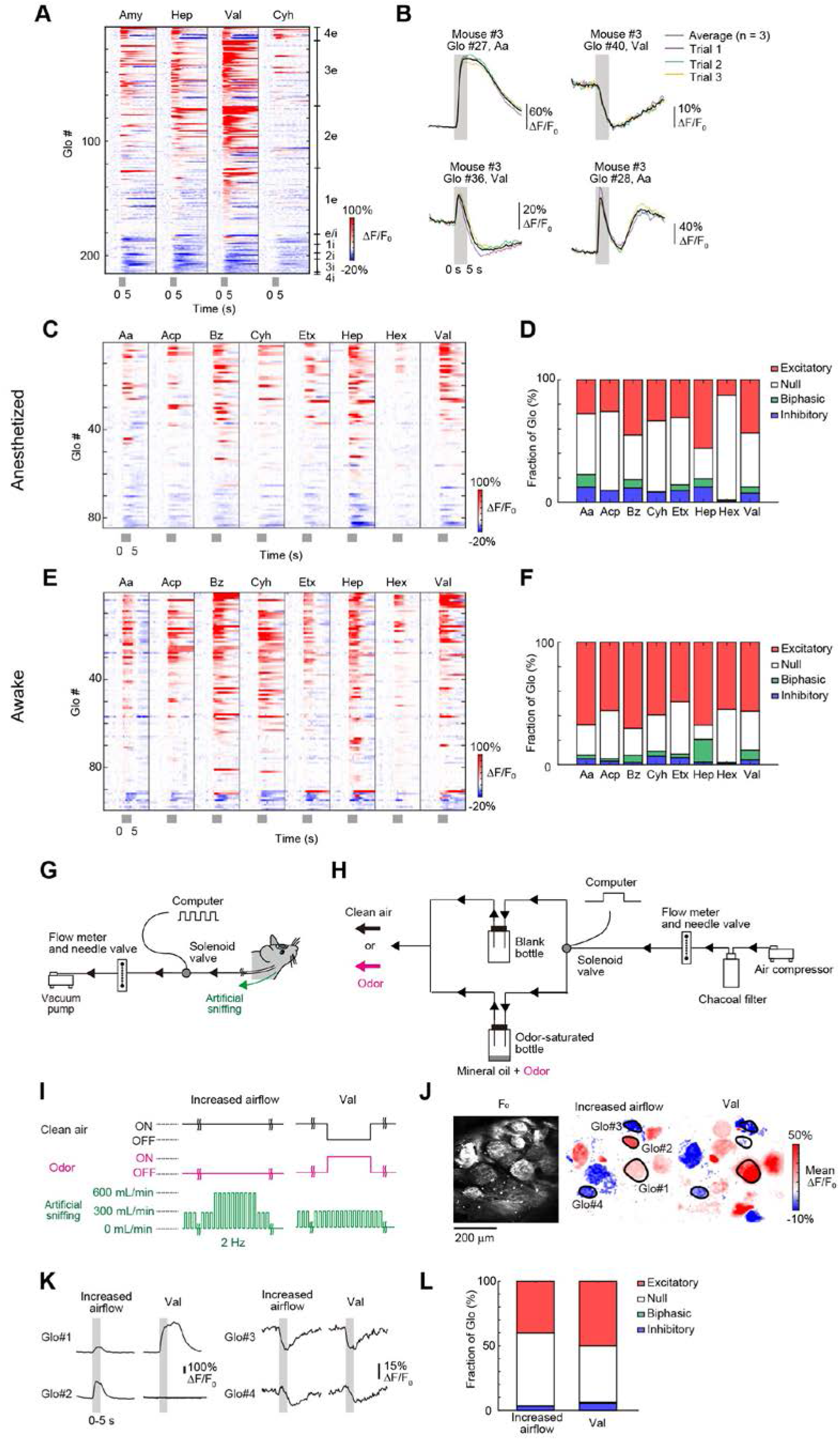
Inhibitory responses at OSN axon terminals, Related to Figure 1. (**A**) Temporal profiles of odor-evoked responses at OSN axon terminals. The responses were sorted based on the number of odorants that elicited odor-evoked responses and their amplitudes. Inhibition was commonly found among different odorants. N = 299 glomeruli from 5 mice. 215 glomeruli showing significant responses to at least one odorant are shown. (**B**) Reproducibility of odor-evoked excitatory and inhibitory responses (N = 3 trials each). The glomeruli showed consistent responses to repeated stimuli. (**C-F**) Widespread inhibitory responses in both anesthetized (**C, D**) and awake (**E, F**) animals (N = 3). Larger excitatory responses seen in awake mice may be due to the higher sniffing speed/rate. Temporal profiles (**C, E**) and the polarity of odor-evoked responses (**D, F**) in all glomeruli are shown. N = 105 glomeruli from 5 mice. 84 and 99 glomeruli showing significant responses to at least one odorant are shown for anesthetized and awake states, respectively. (**G**) Artificial sniffing in tracheotomized mice. Airflow timing was controlled by a solenoid valve. (**H**) Olfactometer used in this study. A solenoid valve was used to switch clean air vs. odorized air. Odorants were diluted in mineral oil. Saturated vapor in the bottle was delivered to the animal. (**I**) Artificial sniffing system to examine mechanosensation (left, Increased airflow) and odor responses (right, Val) in tracheotomized and anesthetized mice. Increased airflow or Val stimulation was applied during artificial sniffing (bellow). The timing of airflow generation was controlled by the opening of electromagnetic valves. (**J**) Excitatory and inhibitory responses were seen for increased airflow (middle) and Val stimuli (right). Excitatory and inhibitory responses are shown in red and blue, respectively. Scale bar, 200 μm. (**K**) Representative mechanosensory responses to increased airflow and Val-evoked responses. Pulsed airflow (2 Hz, 250 ms on and 250 ms off) was artificially produced during the imaging session. Increased airflow (300 to 600 mL/min) or Val stimulation was delivered to the same animal for 5 s (shown in gray). (**L**) Polarity of glomerular responses to increased airflow and Val stimuli. N = 149 glomeruli from 3 mice.

**Figure S2.**
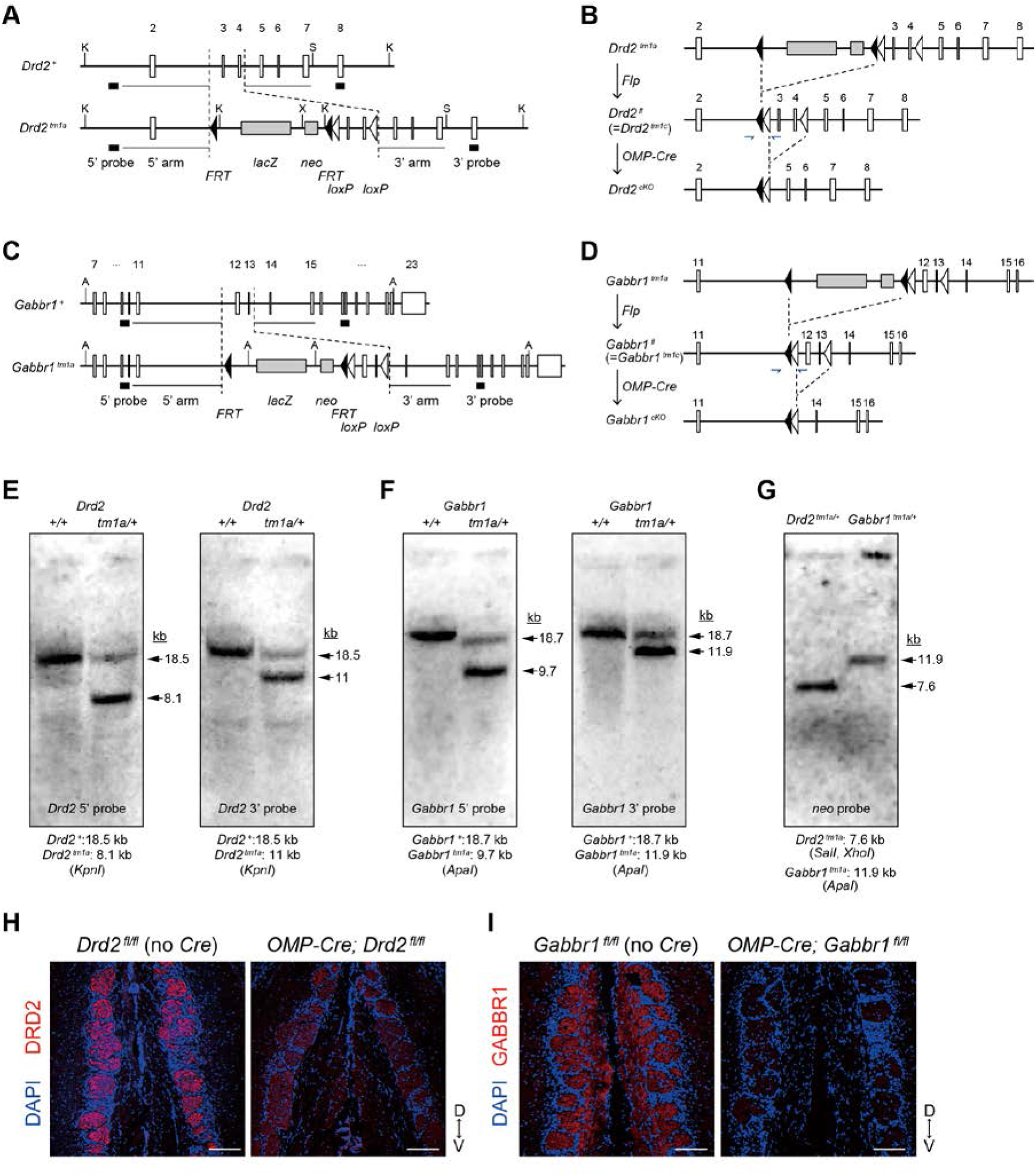
OSN-specific GABAR_B1_ and D2R knockout mice, Related to Figure 2. (A) Gene targeting at *Drd2* locus. We obtained germline transmission from one ES clone, HEPD064_5_D11. The *Drd2* locus in *Drd2*^*tm1a*^, *Drd2*^*fl*^ (= *Drd*^*tm1c*^) and *Drd*^*cKO*^. *Drd2*^+/*tm1a*^ mice were crossed with Flp mice to obtain *Drd2*^+/*fl*^. (**C**) Gene targeting at *Gabbr1* locus. We obtained germline transmission from one ES clone, EPD0730_1_G04. (**D**) The *Gabbr1* locus in *Gabbr1*^*tm1a*^, *Gabbr1*^*fl*^ (=*Gabbr1*^*tm1c*^), and *Gabbr1*^*cKO*^. *Gabbr1*^+/*tm1a*^ mice were crossed with *Flp* mice to obtain *Gabbr1*^+/*fl*^. (**E-G**) Southern blot analysis on *Drd2*^+/+^ and *Drd2*^*tm1a*/+^ (**E**, *Drd2* probe), *Gabbr1*^+/+^ and *Gabbr1*^*tm1a*/+^ (**F**, *Gabbr1* probe); *Drd2*^*tm1a*/+^ and *Gabbr1*^*tm1a*/+^ (**G**, *neo*^*r*^ probe). (**H**) Immunostaining of D2R in the OB of OSN-specific *Drd2* mutant mice. (**I**) Immunostaining of GABAR_B1_ in the OB of OSN-specific *Gabbr1* mutant mice. Location of 5’ and 3’ arms for gene targeting and 5’ and 3’ DNA probes for southern blotting are shown in (**A**) and (**C**). *KpnI* (K), *SpeI* (S), *XhoI* (X) and *ApaI* (A).

**Figure S3.**
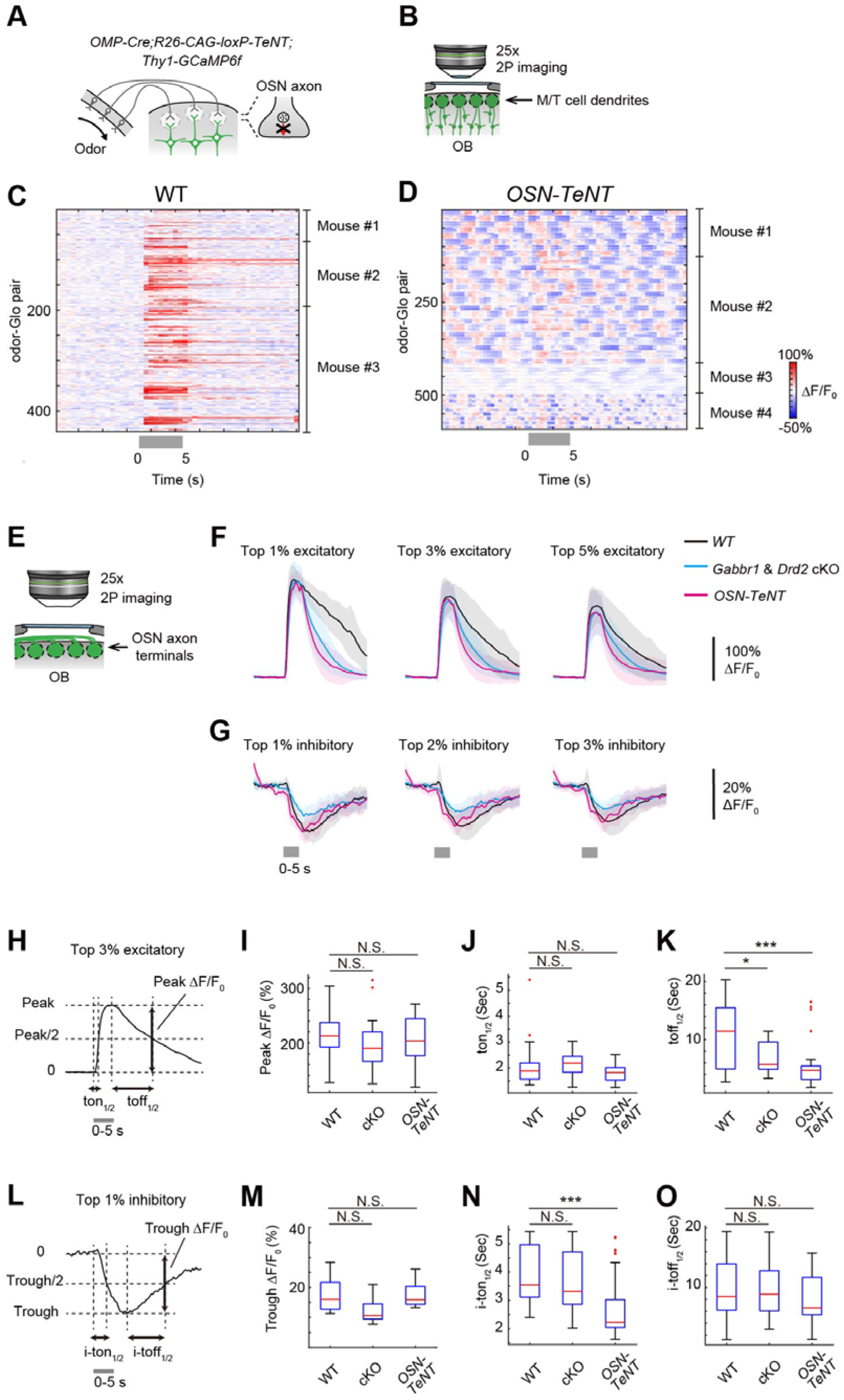
OSN-specific TeNT knock-in mice and analysis of odor-evoked responses in mutant mice, Related to Figure 2. (**A**) *OMP-Cre* knock-in mice were crossed with *R26-CAG-loxP-TeNT* knock-in mice to obtain OSN-specific TeNT knock-in mice (OSN-TeNT). VAMP2 staining was eliminated at OSN axon terminals in this mouse (Fujimoto et al., 2019). (**B**) *Thy1-GCaMP6f* mouse line was used to examine the odor responses of OSN-TeNT mice. (**C, D**) Odor-evoked responses of M/T cells in wild-type (**C**) and OSN-TeNT mice (**D**). Summary of all odor-glomeruli pairs are shown. Odor-evoked responses were almost completely abolished in OSN-TeNT mice. Instead, the M/T cells in all glomeruli demonstrated synchronized spontaneous activity every a few seconds (**D**). This may indicate that OSN input is required for desynchronized spontaneous activity in M/T cells. WT, N = 440 odor-glomerulus pairs (55 glomeruli) from 3 mice; OSN-TeNT, N = 592 odor-glomerulus pairs (74 glomeruli) from 4 mice. Tested odorants are amyl acetate, acetophenone, benzaldehyde, cyclohexanone, ethyl hexanoate, heptanal, hexanoic acid, and valeraldehyde, diluted at 0.5%. (**E-O**) Temporal kinetics of excitatory and inhibitory responses of OSN axon terminals averaged across different fraction of odor-glomerulus pairs. For fair comparison across mutant lines, the top excitatory (1, 3 and 5%; **F**) and top inhibitory (1, 2 and 3%; **G**) fractions (based on the mean ΔF/F_0_) were used to show averaged excitatory and inhibitory response kinetics, respectively. (**H-K**) Temporal profiles of excitatory responses. The rise time (time to half-maximum, ton_1/2_) and decay time (time from peak to half-maximum, toff_1/2_) were compared among wild-type, *Gabbr1/Drd2* cKO, and OSN-TeNT. The decay time (toff_1/2_) was significantly shorter in *Gabbr1/Drd2* cKO, and OSN-TeNT. (**L-O**) Temporal profiles of inhibitory responses. The decay time (time to half of the inhibitory trough, i-ton_1/2_) and recovery time (time from the trough to the half of the trough, i-toff_1/2_) were compared. One-way ANOVA and Tukey-kramer post-hoc tests were used. N.S., not significant; *, p < 0.05; **, p < 0.01; ***, p < 0.001.

**Figure S4.**
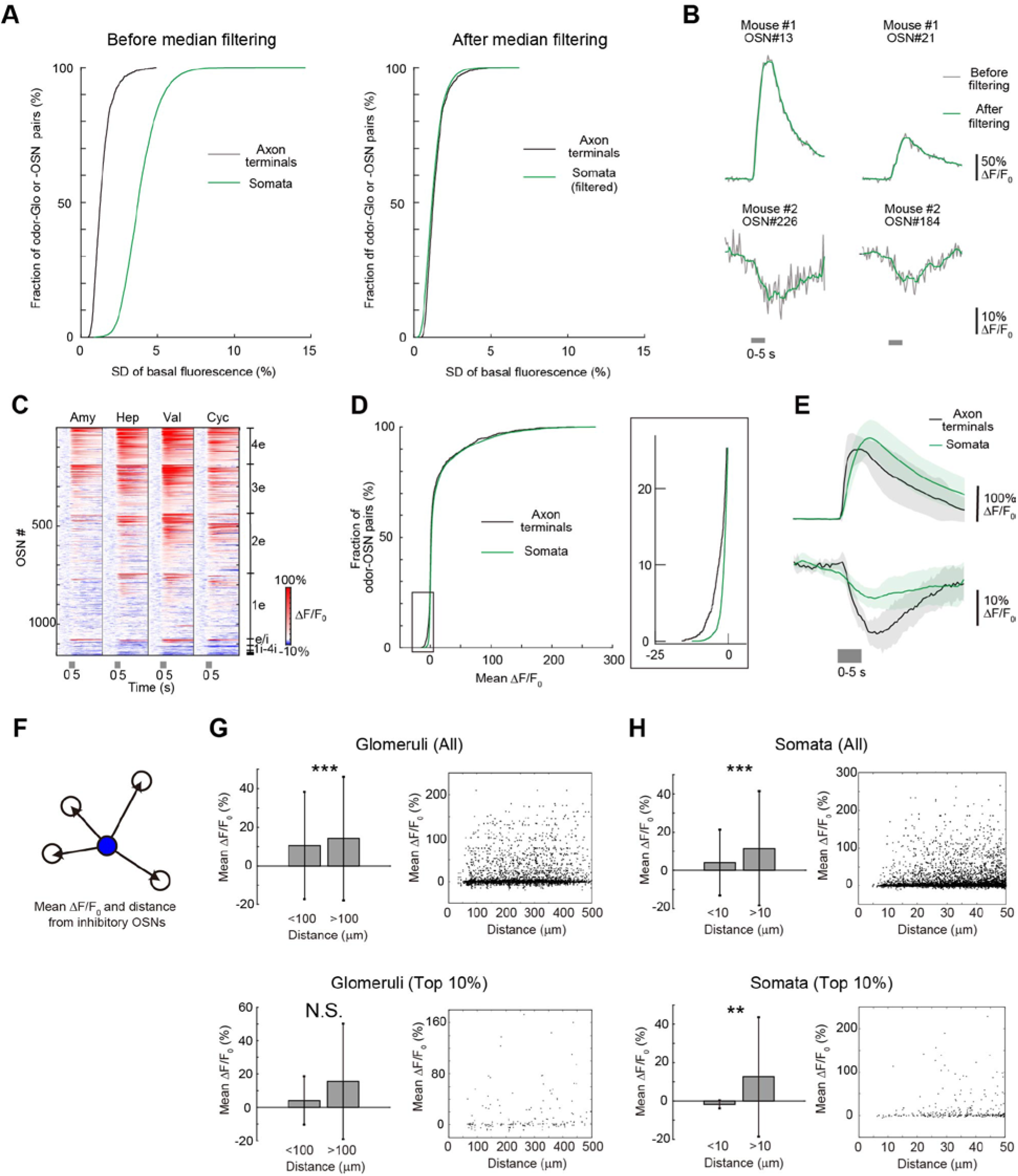
Inhibitory responses in the OSN somata in the OE, Related to Figure 3. (**A**) Basal fluctuation (standard deviation, SD) of GCaMP signals before and after temporal median filtering within a 3-second time window. The SD distribution after filtering (right, green) is almost comparable to that of glomerulus data (right, gray), suggesting a similar noise level. N = 1,196 odor-glomerulus pairs from 5 mice for axon terminals; N = 6,468 odor-OSN pairs from 4 mice for somata. (**B**) Temporal median filtering does not affect the temporal profiles of excitatory and inhibitory responses. In general, inhibitory responses were slower than the excitatory ones, but this is most likely due to the much slower decay of GCaMP3 signals than the rise (Tian et al., 2009). (**C**) Temporal profiles of odor-evoked responses in OSN somata. The responses were sorted by the number of odorants that elicit odor-evoked responses and their amplitude. Inhibition was commonly found for all four odors. N = 1,617 OSNs from 4 mice. 1,159 OSNs showing significant responses to at least one odorant are shown. (**D**) Comparison of OSN axon terminals vs OSN somata. Cumulative histogram of response amplitude for axon terminals and somata. Inset includes expanded *x*- and *y*-axes to display the population showing inhibitory responses. N = 1,196 odor-glomerulus pairs from 5 mice for axon terminals; N = 6,468 odor-OSN pairs from 4 mice for somata. (**E**) Temporal kinetics of excitatory and inhibitory responses in OSN somata and axon terminals. For fair comparisons, the top 3% of excitatory and top 1% of inhibitory OSNs were averaged for excitatory and inhibitory traces, respectively. Data are from 35 and 11 odor-glomerulus pairs for excitatory and inhibitory responses, respectively, at axon terminals; Data are from 194 and 64 odor-OSN pairs for somata. (**F-H**) To investigate the possible contribution of ephaptic coupling in inhibitory responses, we examined odor-evoked responses in glomeruli (**G,** upper, N = 334 and 3,093 glomeruli from 5 mice) and OSNs (**H,** upper, N = 123 and 135,058 OSNs from 4 mice) surrounding inhibitory ones. However, glomeruli located within 100 μm (adjacent) were not more excitatory than others. Similarly, OSNs located within 10 μm (adjacent) were not particularly more excitatory than others. The conclusion remained the same when we only focused on top 10% inhibitory glomeruli/OSNs (**G**, lower, N = 24 and 176 glomeruli; **H**, lower, N = 7 and 5864 OSNs).

**Figure S5.**
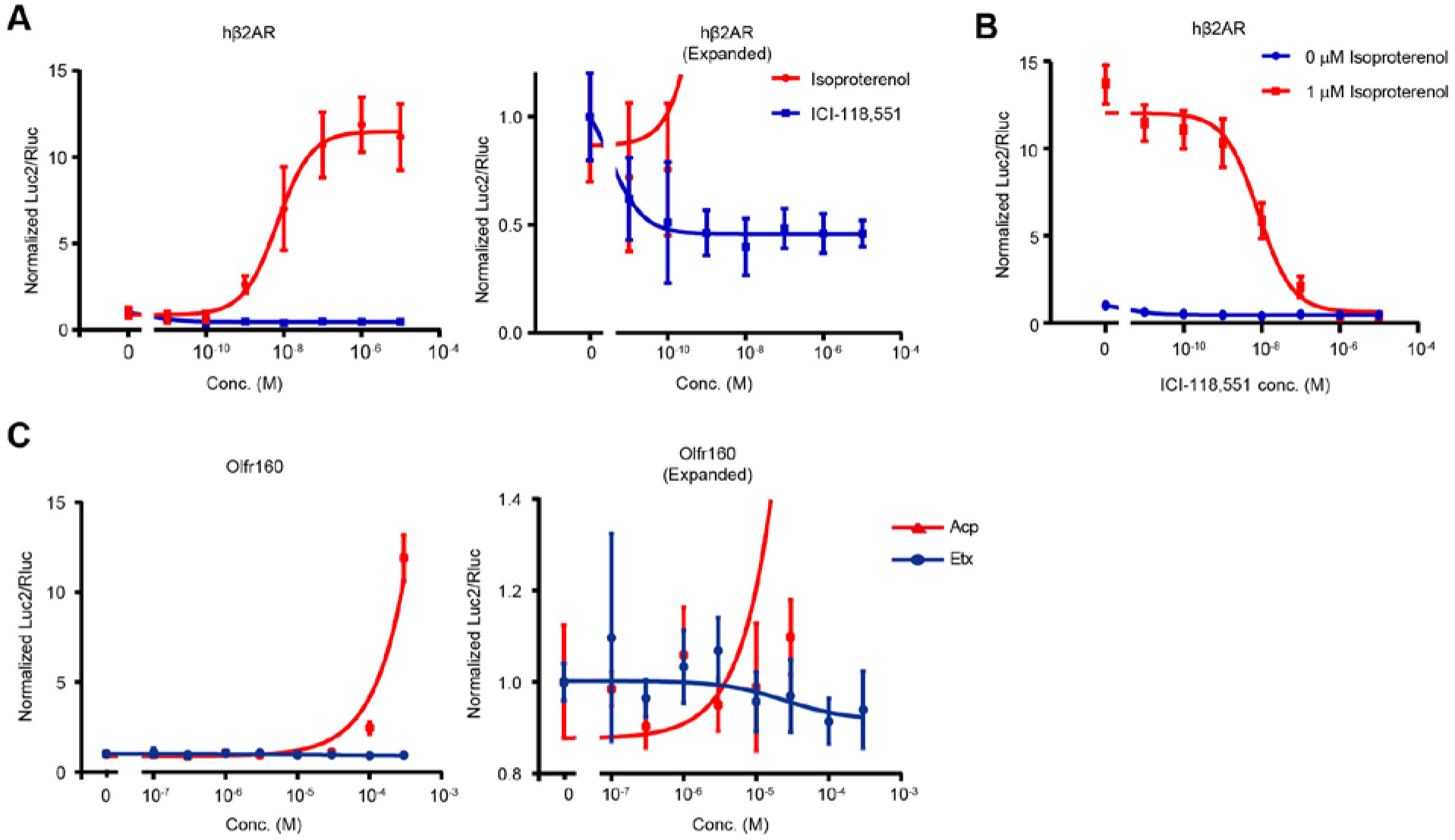
A heterologous assay system for ORs, Related to Figure 5. (**A**) Control experiments for the luciferase assay of CRE activity. Human β2-adrenergic receptor (hβ2AR) also couple to Gs/olf. Here we examined the responses of hβ2AR to an agonist, isoproterenol, and an inverse agonist ICI-118,551 (ICI). ICI demonstrated suppression of basal CRE activity in the luciferase assay. (**B**) ICI also inhibited hβ2AR responses to isoproterenol in a dose-dependent manner. (**C**) Odor responses of Olfr160. Acetophenone acts as an agonist but ethyl hexanoate acts as an inverse agonist toward Olfr160.

**Figure S6.**
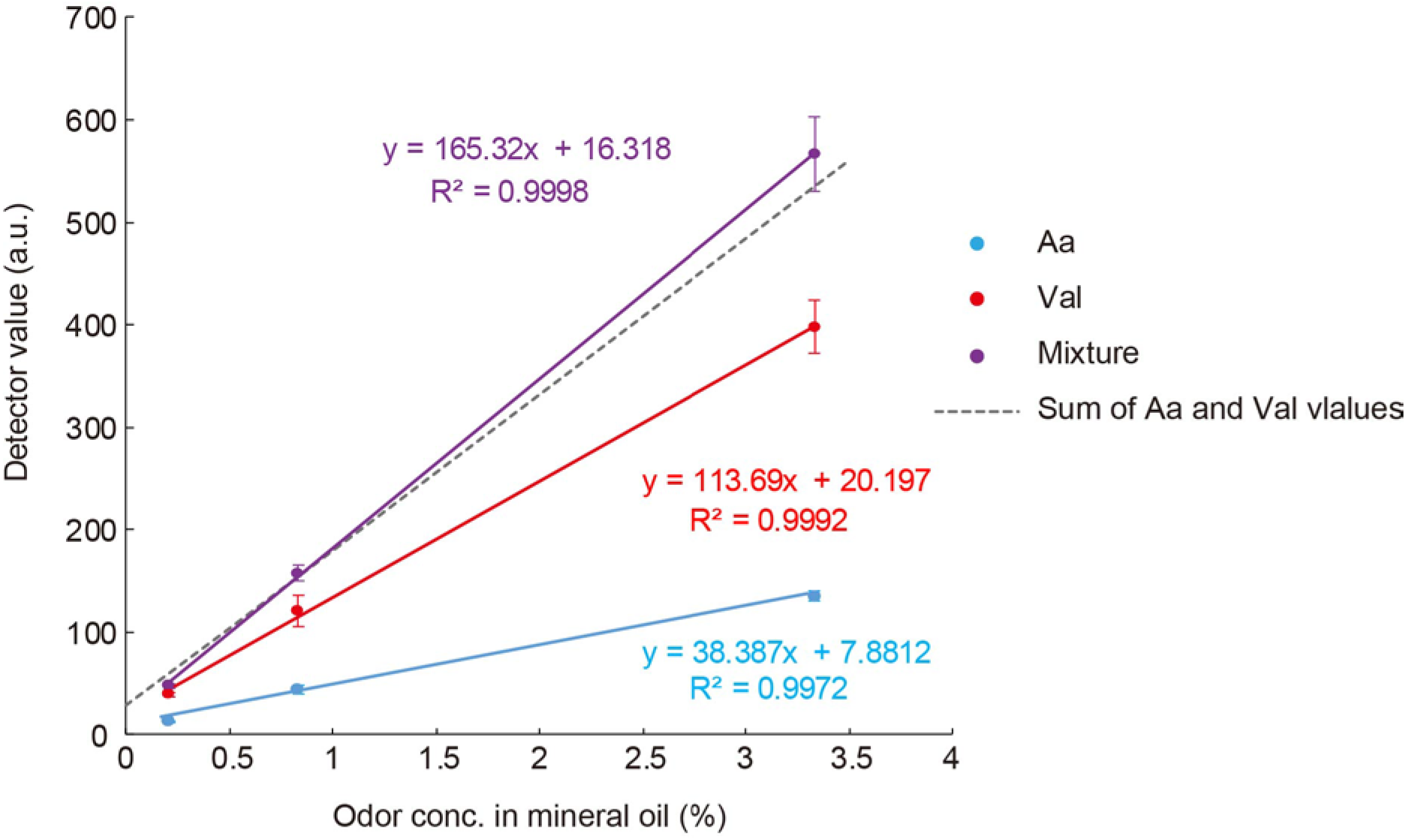
Measurements of odor concentrations in the saturated vapor, Related to Figure 6-8. The relative odor concentration of the saturated vapor in the bottle was measured by a semiconductor-based odor sensor. The air was odorized by amyl acetate (Aa, cyan) and valeraldehyde (Val, red) diluted in mineral oil at 1:480, 1:120, and 1:30 (v/v). Odor mixture (purple) was made by mixing Aa and Val in mineral oil at a defined concentration. The detected value was linearly correlated with the odor concentrations in the mineral oil in the bottle. We also confirmed that the values of odor mixtures (Aa + Val) were linear sum of the values for each component. Sum of Aa and Val values (dotted line) were drawn based on linear sum of the equation for Aa and Val. N = 5 trials in each point. Error bars indicate mean ± standard deviation.

**Figure S7.**
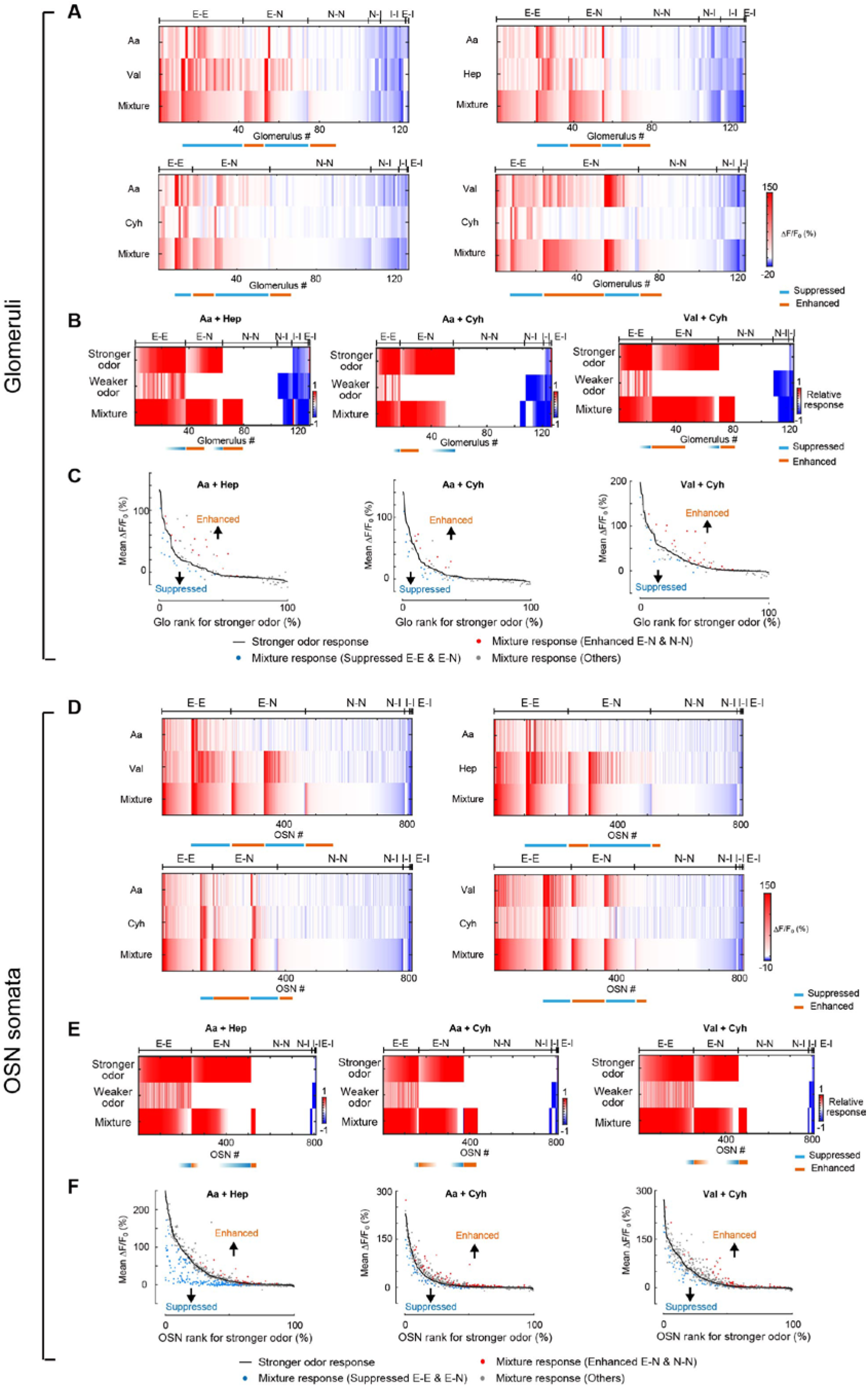
Representation of odor mixtures at OSN axon terminals and OSN somata, Related to Figure 6-8. (**A**) Summary of glomerular responses at axon terminals to different odor mixture pairs (Aa+Val, Aa + Hep, Aa + Cyh, and Val + Cyh) is shown. Excitatory (E) and inhibitory (I) responses were defined when response mean > 3 SD above baseline. All other glomeruli were defined as null (N) response. Glomeruli were categorized based into E-E, E-N, N-N, N-I, I-I, and E-I types. All glomeruli are shown including ones showing null responses. Glomeruli were sorted based on suppression/enhancement and the magnitude of mixture responses. N = 131 glomeruli from 3 mice. (**B**) Stronger and weaker odors were defined for each glomerulus. The relative response indicates a normalized response based on maximum responses among the three conditions. Responses to weaker odor, stronger odor, and their mixture are shown. Glomeruli are sorted based on the mixture responses, and then stronger odor responses. (**C**) Glomeruli demonstrating suppressed and enhanced responses in mixture experiments. Glomerulus rank was defined based on the response amplitude to the stronger odor. Suppression seen for E-E and E-N glomeruli are shown in blue, and enhancement seen for E-N and N-N glomeruli are shown in red. All other glomeruli are shown in gray. (**D**) Summary of OSN soma responses to different odor mixture pairs (Aa + Val, Aa + Hep, Aa + Cyh, and Val + Cyh) are shown. Excitatory (E) and inhibitory (I) responses were defined when response was mean > 3 SD above baseline. All other OSNs were defined as a null (N) response. OSNs were categorized based into E-E, E-N, N-N, N-I, I-I, and E-I types. All OSNs are shown including ones showing null responses. OSNs were sorted based on suppression/enhancement and the magnitude of mixture responses. N = 825 OSNs from 3 mice. (**E**) Stronger and weaker odors were defined for each OSN. The relative response indicates a normalized response based on maximum responses among the three conditions. Responses to the weaker odor, the stronger odor, and the mixtures are shown. OSNs are sorted based on the mixture responses, and then the stronger odor responses. (**F**) OSNs demonstrating suppressed (antagonism) and enhanced (synergy) responses in mixture experiments. OSN rank was defined based on the response amplitude to the stronger odor. Suppression seen for E-E and E-N OSNs are shown in blue, and enhancement seen for E-N and N-N glomeruli are shown in red. All other glomeruli are shown in gray.

**Figure S8.**
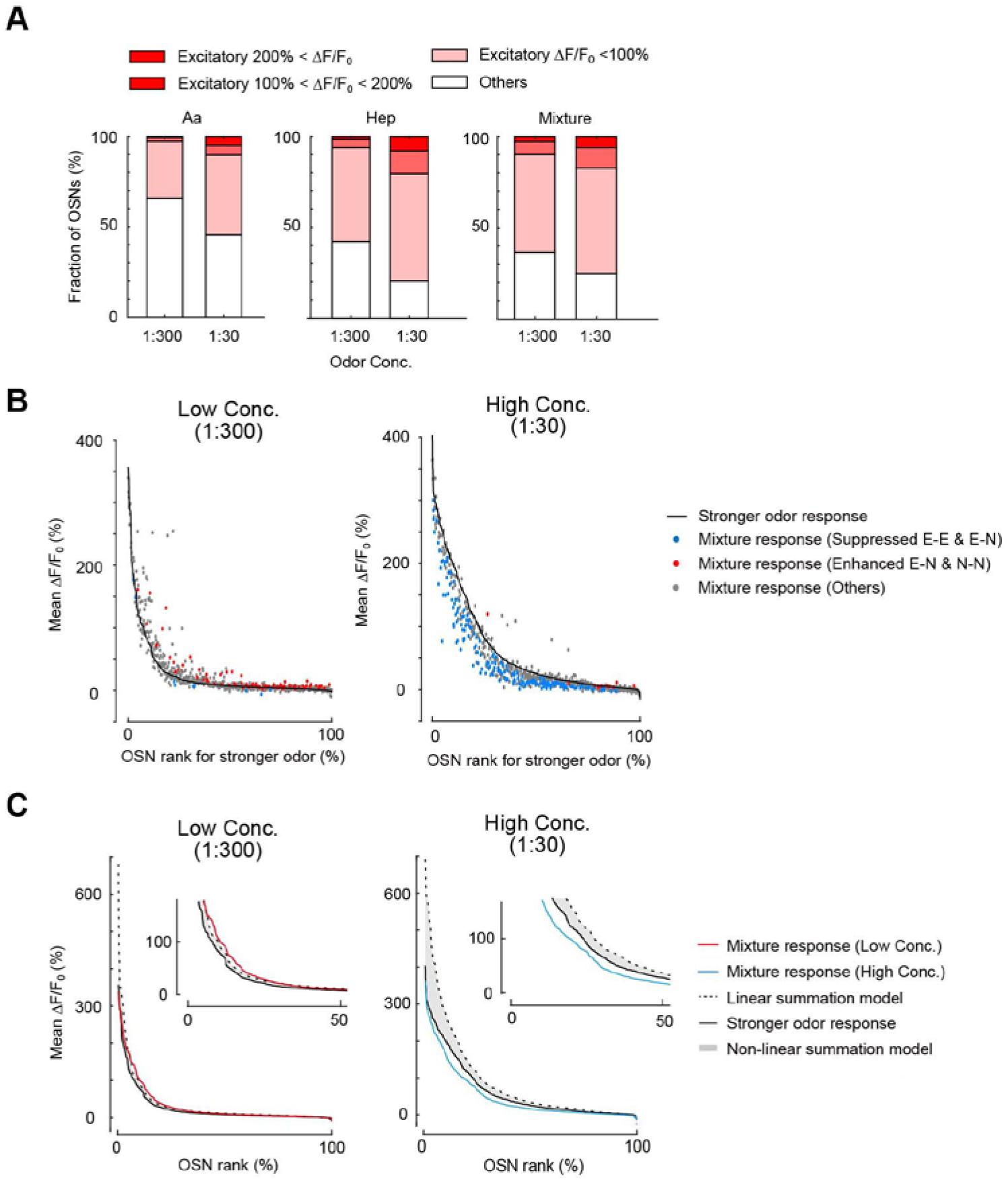
Concentration-dependent mixture effects in OSNs, Related to Figure 8. (**A**) Fractions of OSNs showing excitatory responses to high (1:30) and low (1:300) concentrations of amyl acetate (Aa), heptanal (Hep) and their mixtures. N = 686 OSNs from 3 mice. (**B**) Antagonism and synergy found for high and low concentrations of odor mixtures. OSN rank was defined based on the response amplitude to the stronger odor. Suppression seen for E-E and E-N glomeruli are shown in blue, and enhancement seen for E-N and N-N glomeruli are shown in red. When high concentrations of Aa and Hep were mixed (1:30), suppressed responses (antagonism) were often observed: Suppressed fractions were 25.4% and 1.3% for high and low concentrations, respectively (p < 1.0 × 10^−324^, Chi-square test). In contrast, enhanced responses (synergy) was more prominent when odor concentrations were lower (1:300): Enhanced fractions were 1.8% and 13.2% for high and low concentrations, respectively (p = 7.4 × 10^−11^, Chi-square test). (**C**) Responses to odor mixtures compared to linear and non-linear summation models (shown as gray shade). Predicted and actual responses in OSNs are sorted based on its amplitude. When higher concentrations of odors were mixed, the mixture responses were below the stronger odor curve, indicating that antagonism was dominant in this condition (p = 3.4 × 10^−6^, Kolmogorov-Smirnov test). When lower concentrations of odors were mixed, however, the mixture responses were above the linear summation models, which means that synergy was dominant in this condition. (p = 0.02, Kolmogorov-Smirnov test).

**Movie S1. *In vivo* two-photon imaging of odor responses at the OSN somata in the OE.** *In vivo* two-photon imaging of OSN somata in the OE. An OSN-GCaMP3 mouse was used to image the responses to 0.5% valeraldehyde.

## Notes

### Competing Interest Statement

The authors have declared no competing interest.

### Summary of Updates

Figures 4, 5B, 8, S3H-O, S4A-B, S4F-H, S5A-B, S6, S8: newly added; Figures 1, 2B, 3, 6, 7, S7: revised

